# CenH3-independent kinetochore assembly in Lepidoptera requires CENP-T

**DOI:** 10.1101/836262

**Authors:** N Cortes-Silva, J Ulmer, T Kiuchi, E Hsieh, G Cornilleau, I Ladid, F Dingli, D Loew, S Katsuma, IA Drinnenberg

## Abstract

Accurate chromosome segregation requires assembly of the multiprotein kinetochore complex at centromeres. In most eukaryotes, kinetochore assembly is primed by the histone H3 variant CenH3, which physically interacts with components of the inner kinetochore constitutive-centromere-associated-network (CCAN). Unexpected to its critical function, previous work identified that select eukaryotic lineages, including several insects, have lost CenH3, while having retained homologs of the CCAN. These findings imply alternative CCAN assembly pathways in these organisms that function in CenH3-independent manners. Here, we study the composition and assembly of CenH3-deficient kinetochores of Lepidoptera (butterflies and moths). We show that lepidopteran kinetochores consist of previously identified CCAN homologs as well as additional components including a divergent CENP-T homolog, which are required for accurate mitotic progression. Our study focuses on CENP-T that we find both necessary and sufficient to recruit the Mis12 outer kinetochore complex. In addition, CRISPR-mediated gene editing in *Bombyx mori* establishes an essential function of CENP-T *in vivo*. Finally, the retention of CENP-T homologs in other independently-derived CenH3-deficient insects indicates a conserved mechanism of kinetochore assembly between these lineages. Our study provides the first functional insights into CCAN-based kinetochore assembly pathways that function independently of CenH3, thus contributing to the emerging picture of an unexpected plasticity to build a kinetochore.

## Introduction

The centromere is an essential chromosomal region which ensures equal partitioning of chromosomal DNA during cell division [1]. In all eukaryotes, faithful chromosome segregation requires each chromosome to interact accurately with microtubule fibers from the mitotic or meiotic spindle. This interaction is mediated by the kinetochore, a macromolecular protein complex that assembles on centromeric DNA [2]. The centromere-proximal inner kinetochore hosts components of the Constitutive Centromere Associated Network (CCAN), a group of up to 16 different proteins present throughout the cell cycle that create the centromere-kinetochore interface. Upon cell division, the CCAN recruits the centromere distal outer kinetochore complex, which is composed of the KMN network (Knl1, the Mis12 and the Ndc80 complex) [3]. This recruitment is enabled by two CCAN subunits, CENP-C and CENP-T, that physically interact with subunits of the Mis12 and Ndc80 complexes [4–11]. The KMN network then in turn mediates the interaction with spindle microtubules to drive chromosome segregation during cell division [12].

Previous work identified the histone H3 variant CenH3 (also called CENP-A [13,14]) as a core constituent for defining the site of a functional kinetochore in most eukaryotes. This is because CenH3 forms specialized nucleosomes preferentially found at centromeres that are the target sites for kinetochore assembly [15–18]. To do so, CenH3 physically interacts with CCAN components including CENP-C and CENP-N, thereby connecting the kinetochore to chromatin [19–22]. In addition, the attachment of the kinetochore to chromatin can also be facilitated by other CCAN components with DNA-binding activities. In vertebrates and fungi this includes CENP-C, the direct DNA binding partner of CenH3, and the CENP-T/CENP-W complex, two histone-fold proteins that can form a tetramer with the CENP-S/CENP-X histone-fold dimer [23–25].

Unexpected to its conserved and essential function, CenH3 has been lost in a few select lineages. These include the kinetoplastids [26,27] that have evolved an entirely different kinetochore complex and an early diverging fungus with mosaic centromeres characteristic of both regional and point centromeres [28]. In addition, our previous studies also revealed the recurrent loss of CenH3 in multiple insect species [29]. Interestingly, these species have also undergone changes in their centromeric architecture in which each lineage independently transitioned from monocentric chromosomes (where microtubules attach to a single chromosomal region) to holocentric chromosomes (where microtubules attach along the entire length of the chromosome). This discovery suggests that these insect species employ an alternative kinetochore assembly mechanism that occurs chromosome-wide in a CenH3-independent manner. Interestingly, despite the loss of CenH3, computational predictions revealed the presence of several CCAN and KMN homologs in holocentric insects [29]. While those predictions provided some insights into the composition of these CenH3-deficient kinetochores, key aspects of kinetochore function, including how these kinetochores attach to centromeric chromatin and to the outer kinetochore proteins, remained unresolved.

To address these unknowns, we performed proteomic analyses in cell lines from the holocentric Lepidoptera (butterflies and moths). This revealed the presence of several previously unidentified kinetochore components in these insects including a homolog of CENP-T, a core kinetochore component bridging centromeric DNA to the outer kinetochore in vertebrates and fungi. We show that in Lepidoptera, CENP-T and other kinetochore components are essential for chromosome segregation and the recruitment of the KMN network. Furthermore, we show that homologs of CENP-T are also present in other independently derived CenH3-deficient insects, indicating a conserved mechanism of kinetochore assembly between these CenH3-deficient lineages.

## Results

### Identification of kinetochore components including CENP-T in CenH3-deficient Lepidoptera

To gain insights into the composition of CenH3-independent kinetochores, we performed immunoprecipitation (IP) experiments of kinetochore components that we previously identified in our computational survey [29] (Figure 1A). For this, we established several stable *Spodoptera frugiperda* (Sf9) cell lines expressing 3xFLAG-tagged CCAN (CENP-M, CENP-N, CENP-I) and outer kinetochore (Dsn1 and Nnf1) components to map their protein interaction profiles by mass spectrometry.

**Figure 1:**
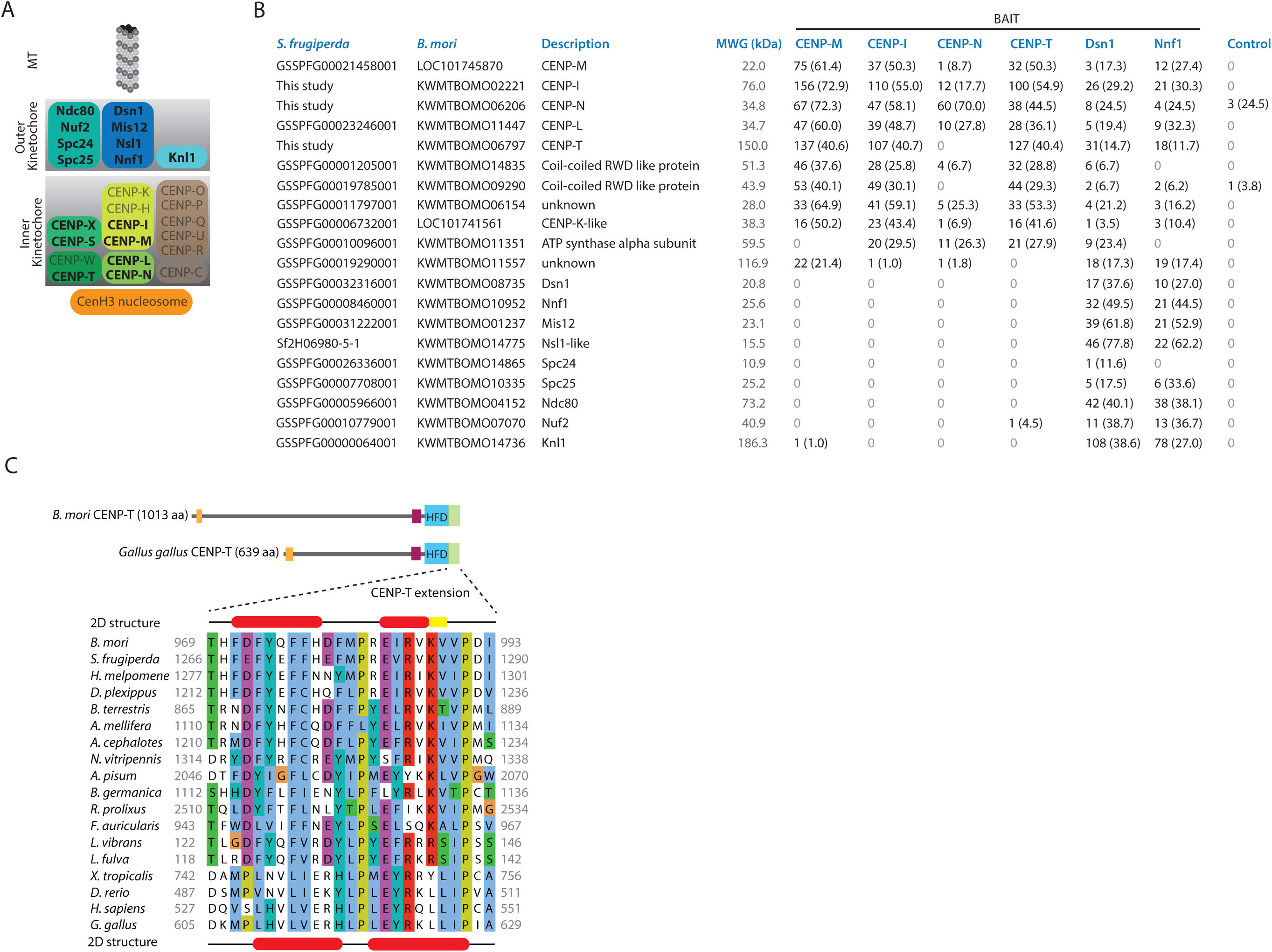
Identification of kinetochore components including CENP-T in CenH3-deficient Lepidoptera. A) Schematic kinetochore organization in vertebrates and fungi. Boxes indicate subcomplexes within inner and outer kinetochores. Kinetochore components present in Lepidoptera are highlighted in bold ([29], this study). B) The table lists the number of peptides and coverages (in parenthesis) of *S. frugiperda* proteins identified by mass spectrometry, which were enriched in the kinetochore IPs over the control. The corresponding homologs in *B. mori* and descriptions based on homology predictions are listed alongside. C) Top: Graphical representation of the *B. mori* CENP-T and *G. gallus* CENP-T sequence features showing the location of the histone fold domains (blue box), CENP-T family specific extension (green box), Arginine-rich (*E* value 4.7×10^−5^ and *E* value 6.0×10^−7^, respectively) and Proline-rich regions (*E* value 3.7×10^−4^ and *E* value 3.4×10^−8^, respectively). Bottom: Multiple alignment of the Cterminal extension of vertebrate and insect CENP-T proteins. *B. mori* (above) and *G. gallus* (below) secondary-structure predictions are derived from the Jpred4 (α-helical regions are in red; β-strands in yellow) [75]. Background colouring of the residues is based on the clustalx colouring scheme. Full-length insect CENP-T sequences and accession numbers if available are listed in Table S2.

Mass spectrometry analyses of the kinetochore immunoprecipitates recovered all of the previously predicted homologs of outer kinetochore and inner kinetochore components, confirming protein complex formation even in organisms that have lost CenH3 (Figure 1B, Figure S1 and Table S1). More importantly, our IPs also revealed several additional components, some of which harbor remote homology to other known kinetochore components (Figure 1B). Among those, we identified a CENP-K-like protein thereby experimentally supporting previous homology predictions in *Bombyx mori* [30]. We also identified two components (GSSPFG00019785001 and GSSPFG00001205001) with remote similarities to the coiled-coil and RWD domains of the budding yeast CENP-O and CENP-P proteins, respectively. Furthermore, we identified the homolog of the alpha subunit of the ATP synthase (bellwether) (GSSPFG00010096001), previously found to localize to kinetochores during *Drosophila melanogaster* male meiosis [31], indicating that this function might be extended to lepidopteran mitosis.

Both inner and outer kinetochore immunoprecipitates also revealed the presence of a 191.2 kDa protein in *S. frugiperda* (GSSPFG00025035001) with KWMTBOMO06797 being the *B. mori* homolog (Figure 1B). Iterative hidden Markov model (HMM) profile searches within annotated proteomes first revealed homologs in several other insect orders and then known vertebrate CENP-T homologs as best hits (Figure S2A). Similarly, HMM profile searches within known protein structures identified the known *Gallus gallus* CENP-T as the best hit (Figure S2B). The alignment between the lepidopteran and vertebrate CENP-T proteins was restricted to their C-terminal parts that include the histone fold domain (HFD). Importantly, the alignment extended to the known 2-helix CENP-T extension with several conserved amino acids being identical, supporting homology to CENP-T rather than to any other histone fold protein (Figure 1C, Figure S2C). The sequence aligning to the CENP-T extension appears to be even better conserved than the HFD allowing for the identification of the *G. gallus* CENP-T in reciprocal HMM profile searches (Figure S2B). Furthermore, analyses of the primary amino acid sequence revealed additional similarities between the *G. gallus* CENP-T and the *B. mori* protein including an enrichment of positively charged amino acids at the N-terminus and a proline patch N-terminal of the HFD (Figure 1C). Finally, reciprocal IP experiment of the *S. frugiperda* homolog confirmed its interaction with kinetochore components (Figure 1B). Although orthology between the lepidopteran proteins and CENP-T cannot be confirmed (Figure S3), the common sequence architecture, kinetochore protein interaction and role in outer kinetochore recruitment (see below) lead us to refer to this lepidopteran component as CENP-T.

Interestingly, we could not detect a putative homolog of the CENP-T histone fold binding partner CENP-W in any of our IP experiments including immunoprecipitates of a 3xFLAG-tagged C-terminal truncated version of CENP-T that contains the HFD and 2-helix extension (Figure S4). Our ability to identify all other known kinetochore homologs in our kinetochore IPs supports the idea that the inability to identify a lepidopteran CENP-W homolog is not due to our IP protocol. However, limitations in detecting those potentially small, arginine-rich proteins by mass spectrometry cannot be excluded.

Taken together, we have identified previously unknown kinetochore components in the CenH3-deficient kinetochore of Lepidoptera. This includes putative homologs of known CCAN components including CENP-T. These results reinforce and extend our previous computational predictions indicating similar kinetochore components in Lepidoptera as those in vertebrates and fungi, despite the loss of CenH3 in the former. Among those, the identification of CENP-T was of particular interest due to its known function in DNA binding and outer kinetochore recruitment in other organisms.

### CENP-T and other kinetochore components are required for accurate mitotic progression

In other organisms, depletion of kinetochore components results in mitotic defects [32]. Previous studies in *B. mori* have shown that, consistently, depletion of outer kinetochore components also results in mitotic defects and increased numbers of aneuploid cells [33]. We extended this previous study by using RNAi to deplete a broader catalogue of kinetochore components. Namely, we depleted several outer kinetochore components of the Mis12 complex (Dsn1, Mis12, Nsl1) and Ndc80 complex (Spc24 and Spc25) as well as several inner kinetochore components (CENP-T, CENP-I, CENP-M, CENP-N). We characterized the resulting mitotic phenotypes by immunofluorescence (IF) at two time points (three and five days). We further validated the efficiency of our kinetochore mRNA depletions by RNA blot analyses (Figure S5).

RNAi-mediated depletion of all kinetochore proteins tested resulted in an increase in the number of mitotic cells relative to the control (RNAi targeting GFP), indicating that kinetochore-depleted cells are delayed or arrested in mitosis. We observed the strongest increase of mitotic cells upon CENP-T, CENP-I and outer kinetochore depletions while milder enrichment was seen upon CENP-M and CENP-N depletion (Figure 2B). Interestingly, while still above control levels, the mitotic indices upon CENP-I, Spc24 and Spc25 depletions were strongly decreased at a later time point (Figure S6A), suggesting that possibly cells can exit from mitosis and continue progressing through the cell cycle or undergo apoptosis.

**Figure 2:**
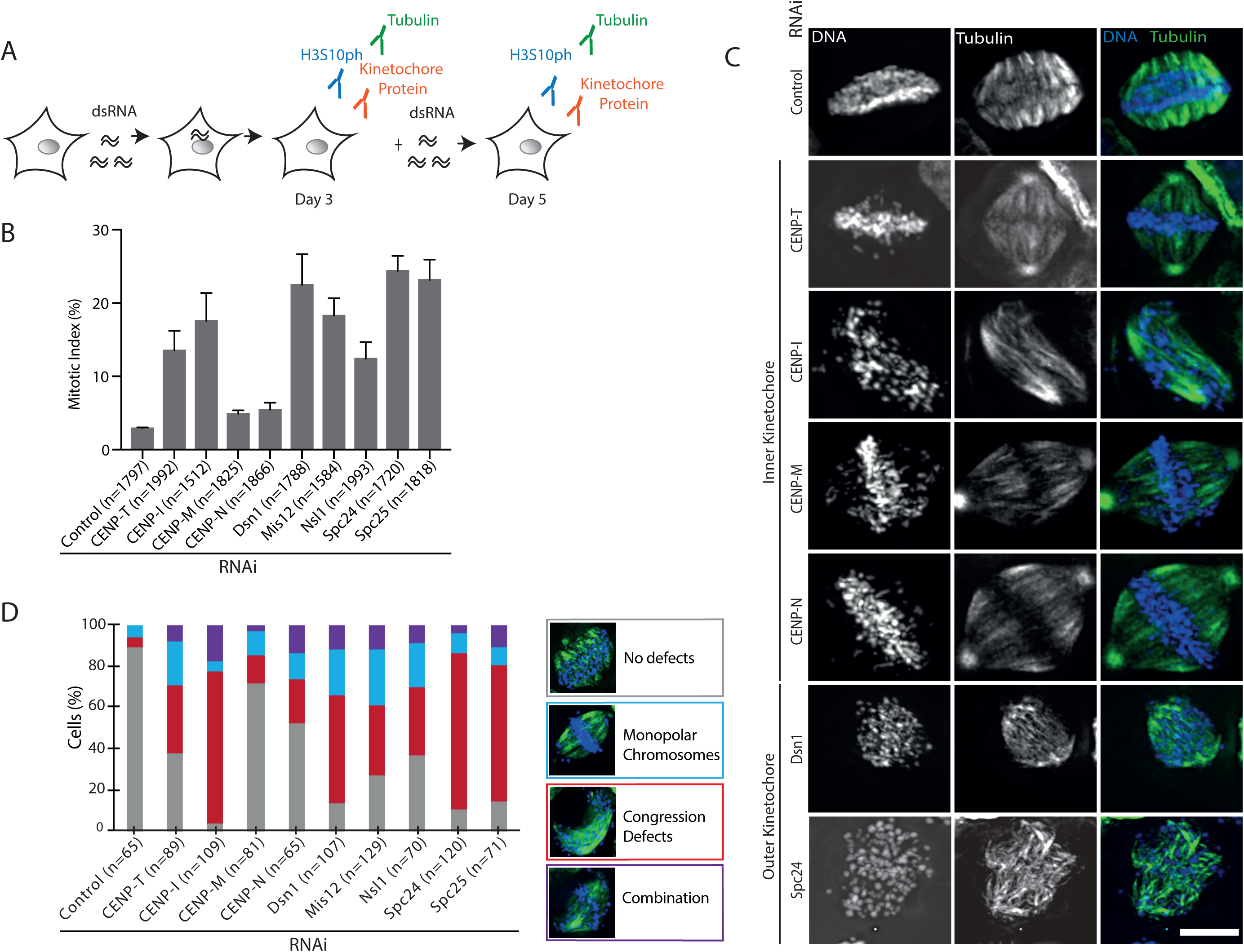
Depletion of kinetochore components affects mitotic progression in *B. mori* cells. A) Schematic of RNAi-mediated depletion strategy. BmN4-SID1 [64] cell lines engineered for properties of enhanced uptake of dsRNA were incubated with various dsRNA constructs for targeted depletion of kinetochore transcripts. After three and five days, cells were analyzed by IF staining against anti-tubulin and anti-phospho Histone H3-Ser10 (H3S10ph) in combination with each one of our custom-made antibodies against the kinetochore protein of interest. Growth medium was changed at day three to add new dsRNA. B) Graph showing the percentage of mitotic cells (H3S10ph positive cells) three days after RNAi-mediated depletions of targeted kinetochore components (n= number of cells, ± standard error of the mean). For results at day five see Figure S6A. C) Representative images of mitotic cells stained with anti-tubulin (green) used to classify mitotic defects observed at three days after depletion of select kinetochore components. Scale bar: 10μm. For zoomed out version see Figure S7. D) Percentage of cells showing no defects (grey), monopolar chromosomes (blue), congression defects (red) and both defects (combination, purple) (n= number of cells analyzed per condition) for different kinetochore depletion experiments.

In addition to the elevated mitotic indices, depletion of kinetochore components results in various degrees of mitotic defects in these cells (Figure 2C, 2D, and Figure S6B, S7, S8). Here, approximately one fourth of CENP-T depleted mitotic cells displayed misaligned, monopolar chromosomes (where one or several chromosomes are not aligned at the metaphase plate and remain at the spindle pole). In addition, 34% of CENP-T-depleted mitotic cells showed complete failure of chromosomes to congress at the metaphase plate (Figure 2C and 2D). This phenotype was even more pronounced upon CENP-I, Spc24 and Spc25 depletion, with the majority of cells (73%, 76% or 66%, respectively) unable to successfully congress chromosomes (Figure 2C and 2D). In contrast, mitotic defects upon CENP-M and CENP-N depletion (Figure 2C and 2D) were relatively mild but became more pronounced at the later time point (Figure S7 and S8). These results show that all tested inner and outer kinetochore components are required for high fidelity chromosome segregation in *B. mori* cells. Additionally, the defects observed upon depletion of the newly identified CENP-T further support its role in chromosome segregation, which we aimed to analyze further.

### The CENP-T N- and HFD containing C-terminus are both essential for accurate mitoses

We next tested whether we can rescue CENP-T knock-down phenotypes by complementation with a RNAi-resistant version of CENP-T. Using double-stranded RNA (dsRNA) targeting the endogenous CENP-T transcript and a FLAG-tagged recoded CENP-T construct unable to be targeted by the dsRNA, we were able to selectively deplete *B. mori* cells of endogenous CENP-T but not FLAG-tagged CENP-T (Figure S9). These experiments were more qualitative than quantitative due to the low transfection efficiency in lepidopteran cells in combination with low numbers of mitotic cells that can be analyzed. Still, among the cells that could be analyzed, expressing the full-length RNAi-resistant construct (CENP-Tres-FLAG) rescued the mitotic defects described above (Figure S9). In contrast, cells expressing the wild-type, non-resistant construct (CENP-T-FLAG) displayed mitotic defects as previously observed (Figure 2). To evaluate whether the N- or HFD containing C-terminus of the CENP-T protein, or both, are required for its function, we expressed RNAi-resistant FLAG-tagged N- and C-terminal truncated CENP-T constructs (ΔN-CENP-Tres-FLAG and ΔC-CENP-Tres-FLAG). Cells expressing either of the two constructs failed to rescue the observed mitotic defects (Figure S9), leading to the conclusion that the full-length CENP-T protein is necessary for accurate mitosis.

### Accurate localization of CENP-T is dependent on other inner kinetochore components and is necessary to recruit the Mis12 complex

Next, to evaluate the role of CCAN and outer kinetochore components in kinetochore assembly we depleted individual kinetochore proteins and stained with custom made antibodies against the lepidopteran CENP-T, Dsn1 (Mis12 complex) and the Spc24/25 (Ndc80 complex). We validated the specificity of the antibodies by mass spectrometry, western blot analyses and IF upon RNAi-mediated depletion of the respective genes (Figure S10).

As expected for kinetochore components, we observed the immunosignals of CENP-T, Dsn1 and Spc24/25 co-localizing with mitotic chromosomes (H3S10ph positive cells) in control BmN4 cells (RNAi targeting GFP) (Figure 3A). We then quantified the intensity of the immunosignals on mitotic chromosomes in control and kinetochore depleted cells (Figure 3B). Depletion of CENP-I, CENP-M and CENP-N significantly impaired CENP-T localization to mitotic chromatin. In contrast, in cells depleted for outer kinetochore components (Dsn1, Mis12, Nsl1 and Spc24) the CENP-T immunosignal could still be observed (Figure 3B and 3C). In addition, depletion of any CCAN component resulted in the loss of Dsn1. In contrast, the recruitment of Spc24/25, while reduced upon other CCAN depletion, was only completely abolished upon depletion of CENP-I (Figure 3B and 3C), consistent with its severe chromosome alignment defects (Figure 2). Dsn1 and Spc24/25 localization was abolished in cells depleted of other members of their respective complexes (Figure 3B and 3C, Figure S11). These data suggest that in *B. mori*, the recruitment of CENP-T is dependent on other inner kinetochore components. In turn, CENP-T appears to be required to recruit the Mis12 but not the Ndc80 complex.

**Figure 3:**
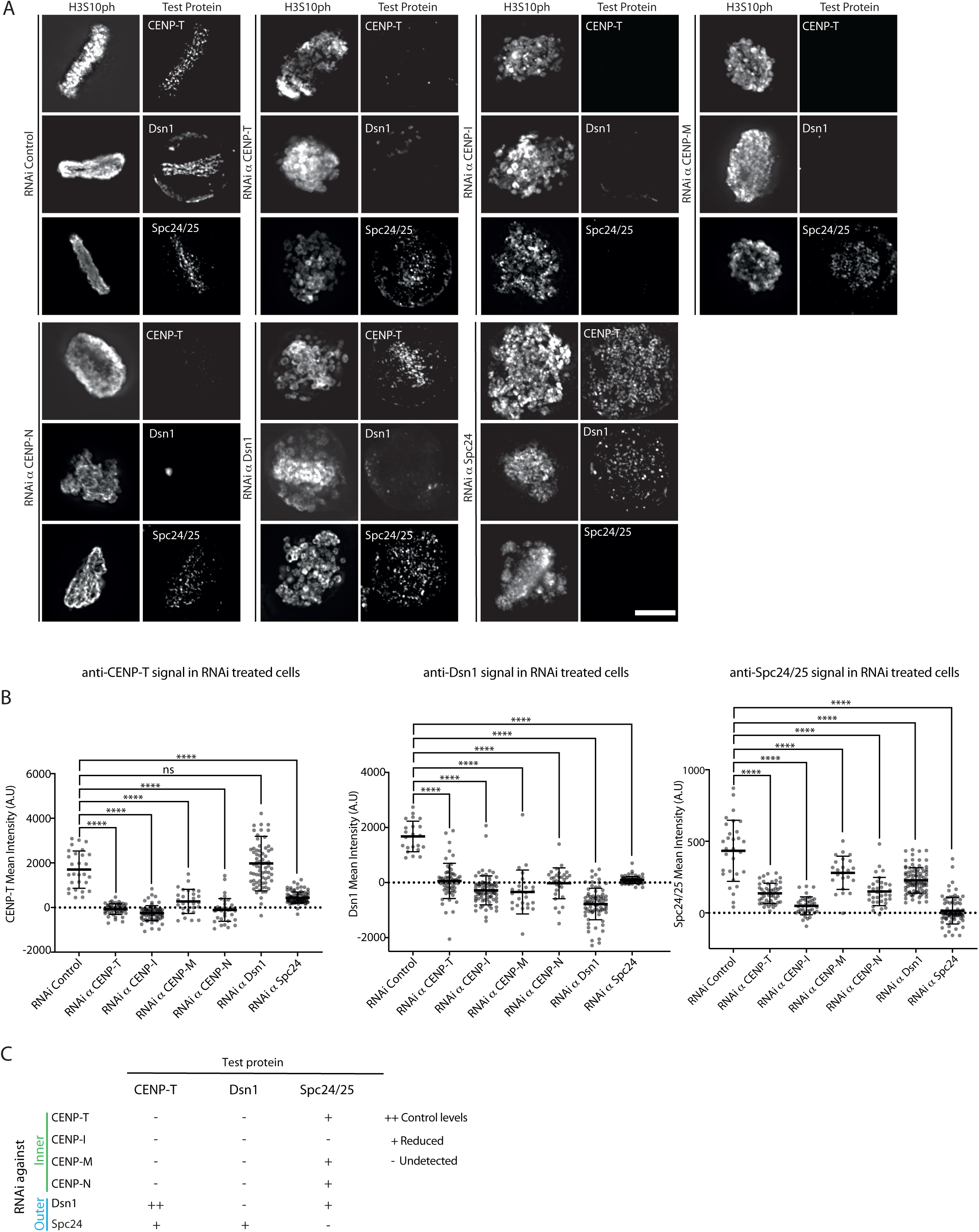
Depletion of CCAN components affects CENP-T and outer kinetochore recruitment in *B. mori* cells. A) Representative images of mitotic BmN4-SID1 cells showing the levels of endogenous CENP-T, Dsn1 and Spc24/25 with and without depletion of inner (CENP-T, CENP-I, CENP-M, CENP-N) and outer (Dsn1 and Spc24) kinetochore components. Scale bar: 10μm. B) Quantification of mean fluorescence intensity for CENP-T, Dsn1 and Spc24/25 signals in the control or upon kinetochore depletions. Statistical significance was tested using the Mann and Whitney test (p< 0.0001, ns= not significant). C) Table summarizing the centromeric localization results of endogenous CENP-T, Dsn1 and Spc24/25 upon RNAi depletions. ++ represents control levels, + represents reduced fluorescence levels compared to control levels and – refers to undetected levels.

### Targeting CENP-T to ectopic sites is sufficient to recruit endogenous CENP-T and the Mis12 complex

Having shown that the inner kinetochore components including CENP-T are necessary for Mis12 complex recruitment, our next aim was to evaluate whether CENP-T is sufficient to recruit outer kinetochore components. For this, we switched to Sf9 cells and developed a stable polyclonal cell line containing Lac operator (LacO) arrays, which enables the targeting of transiently expressed CENP-T-GFP-LacI protein or control (LacI-GFP) constructs to these loci.

In interphase, the ectopic CENP-T-GFP-LacI protein formed circular foci in the nucleus, while in metaphase, CENP-T-GFP-LacI foci resembled filament-like structures. The changes in foci shape suggested a force being applied to these loci in mitosis. In contrast, LacI-GFP foci remained circular in both interphase and mitosis (Figure 4). We then tested whether ectopic CENP-T-GFP-LacI is capable of recruiting endogenous kinetochore proteins using our custom-made antibodies. As expected the anti-CENP-T immunosignal colocalizes with the GFP signal confirming full-length CENP-T protein recruitment to the LacO array. In addition, the anti-Dsn1, but not the anti-Spc24/25 immunosignals colocalized with CENP-T-GFP-LacI foci demonstrating that ectopic CENP-T can recruit endogenous Dsn1. In contrast, in cells transiently expressing LacI-GFP alone, as expected, neither CENP-T, Dsn1 nor Spc24/25 immunosignals colocalized with the ectopic LacI-GFP foci (Figure 4).

**Figure 4:**
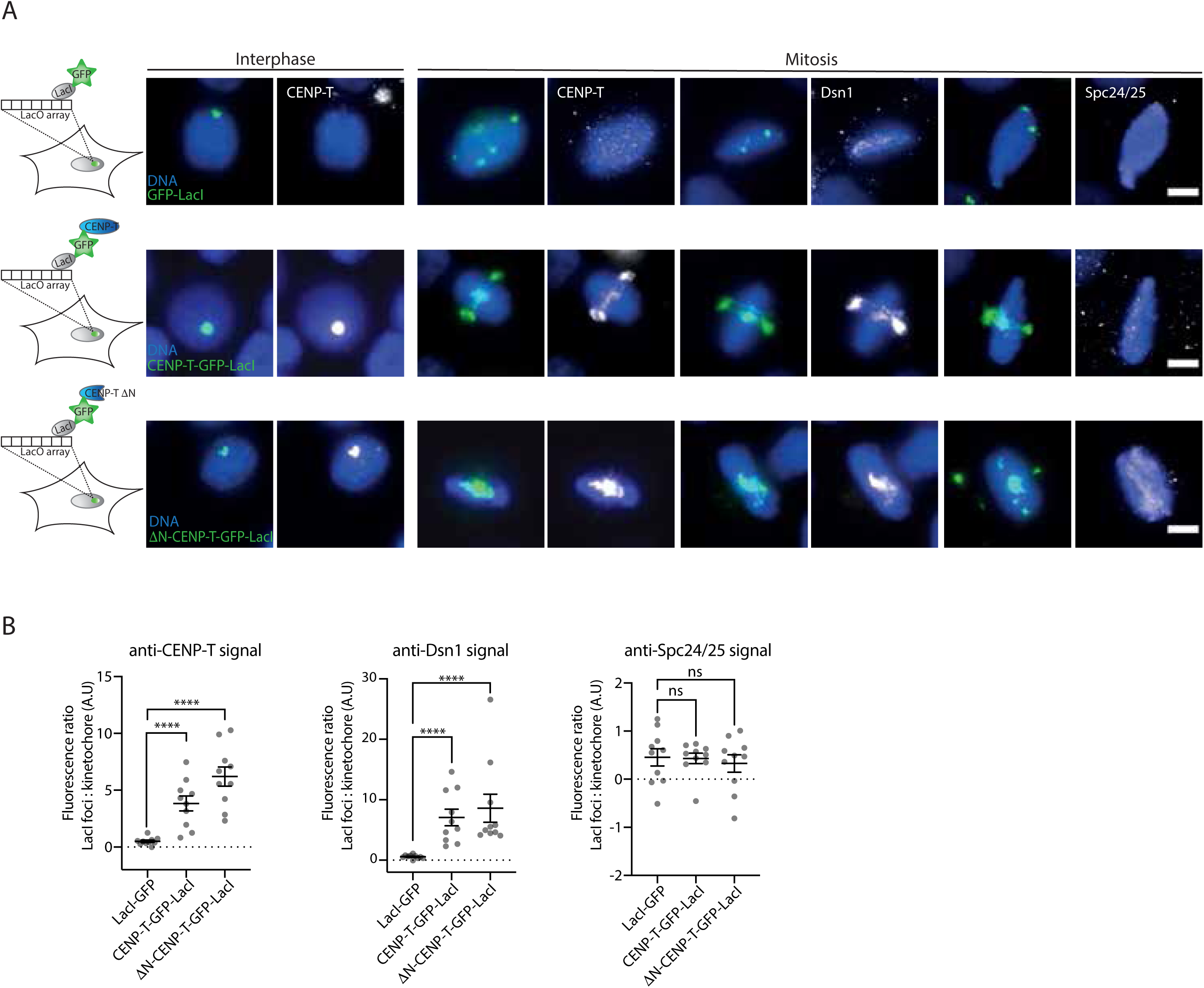
Ectopic CENP-T and CENP-I are sufficient to recruit the outer kinetochore in *S. frugiperda* cells. A) Representative images showing localization pattern of endogenous CENP-T, Dsn1 and Spc24/25 (in grey) in Sf9-LacO cells transiently expressing LacI-GFP (top row), CENP-T-GFP-LacI (second row) and ΔN-CENP-T-GFP-LacI (bottom row) constructs (in green). Endogenous CENP-T and Dsn1 (in grey) colocalize with CENP-T-GFP-LacI and ΔN-CENP-T-GFP-LacI, but not with LacI-GFP foci. Scale bar: 5μm. B) Quantifications of mean fluorescence intensity ratio of CENP-T, Dsn1 and Spc24/25 at the LacI foci compared to levels of CENP-T, Dsn1 and Spc24/25, respectively at endogenous sites (n= 10 cells analyzed). Statistical significance was tested using the Mann and Whitney test (p< 0.0001)

To determine whether ectopic CENP-T-GFP-LacI was able to recruit the endogenous CENP-T, we transiently expressed a truncated CENP-T-GFP-LacI protein (ΔN-CENP-T-GFP-LacI) unable to be recognized by our anti-CENP-T antibody raised against the N-terminus of lepidopteran CENP-T. This allowed us to distinguish between the ectopically expressed ΔN-CENP-T-GFP-LacI (undetected by the anti-CENP-T antibody) and the endogenous CENP-T. Similar to cells expressing full-length CENP-T-GFP-LacI, we found that ΔN-CENP-T-GFP-LacI formed a filament-like structures and colocalized with endogenous Dsn1 but not with endogenous Spc24/25. In addition, we also observed CENP-T immunosignal colocalizing with ΔN-CENP-T-GFP-LacI (Figure 4). Taken together, we conclude that CENP-T is not only necessary but also sufficient for the recruitment of endogenous CENP-T and the Mis12 complex component Dsn1.

### CENP-T is essential *in vivo*

We next aimed to evaluate the importance of CENP-T *in vivo*. For this, CRISPR/Cas9-mediated gene editing was applied to introduce mutations into the endogenous CENP-T gene of the *B. mori* N4 reference strain. A mutant (+7) was isolated that contains a 7-base pair insertion within the guide RNA target site causing a pre-mature STOP codon in the CENP-T gene (Figure 5A). Since the C-terminus of *B. mori* CENP-T containing the HFD and CENP-T extension appears to be essential for its function (Figure S9), the premature stop codon will lead to a non-functional protein product. Heterozygous CENP-T mutants can be readily propagated but when crossed to each other the proportion of unhatched eggs increased to about 30% compared to control crosses (Figure 5B). Genotypic analyses of the progeny of this cross coming from 70% eggs that hatched did not reveal any homozygous CENP-T mutants (Figure 5C). These results show that as *in vitro*, CENP-T is also essential for *B. mori in vivo*.

**Figure 5:**
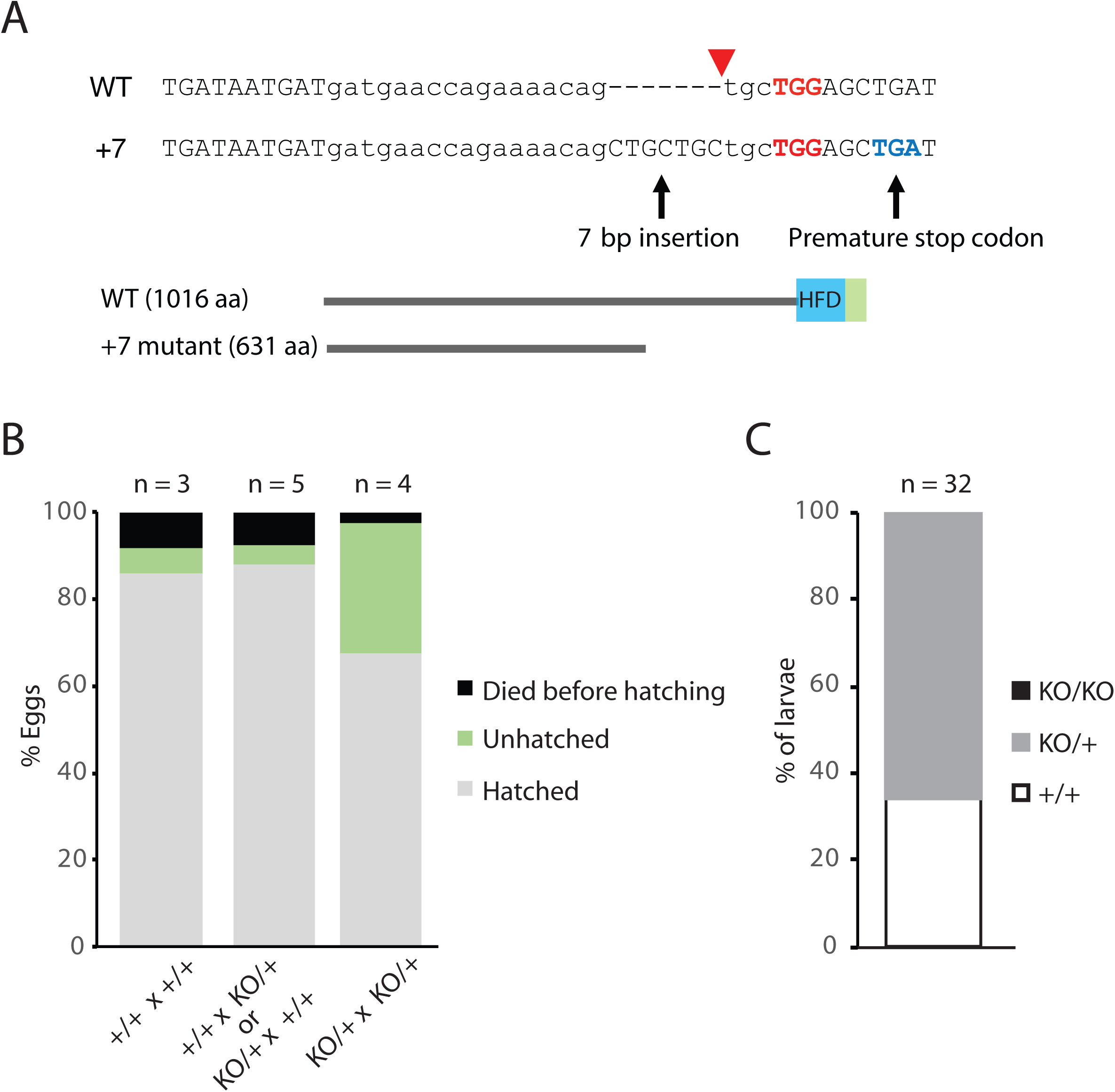
The *B. mori* CENP-T protein is essential *in vivo*. A) CRISPR/Cas9-introduced mutation in the Cenp-T coding sequence. Top: Alignment between wild-type (WT) and mutant (+7) sequences showing a seven base-pair insertion of the Cenp-T gene. Bottom: Schematic of WT and truncated protein products in the CRISPR mutant. PAM site (red), premature stop codon (blue) and Cas9 cleavage site (red arrow) are indicated. B) Hatching rate of the WT and Cenp-T mutant strains. The percentages of hatched (grey), unhatched (green) and eggs that died prior to hatching (black) are indicated for the different crossing patterns between WT (+/+) and heterozygous Cenp-T mutants (+/KO or KO/+). The number of independent crosses n is shown above. The raw data are shown in Table S3. C) Percentages of genotypes from larvae derived from the cross between two heterozygous Cenp-T mutants. The number n of analyzed larvae is indicated. The raw data are shown in Table S4.

### Homologs of CENP-T are retained in independently derived CenH3 deficient insects

Having characterized the function and importance of CENP-T for CenH3-independent kinetochore formation in Lepidoptera, we next profiled its conservation across other CenH3-deficient and CenH3-encoding insects. Orthologs of the lepidopteran CENP-T protein can be readily identified using BLASTP in all CenH3-deficient insects analyzed as well as several CenH3-encoding insects (Figure 6, Table S2). Notably, phylogenetic analyses of HFD proteins belonging to various HFD families including CENP-T indicated accelerated rates of CENP-T protein sequence evolution ancestral to all insects (Figure S3). Importantly, the retention of CENP-T homologs in independently derived CenH3-deficient insects orders indicates important roles for CENP-T in kinetochores in these organisms.

**Figure 6:**
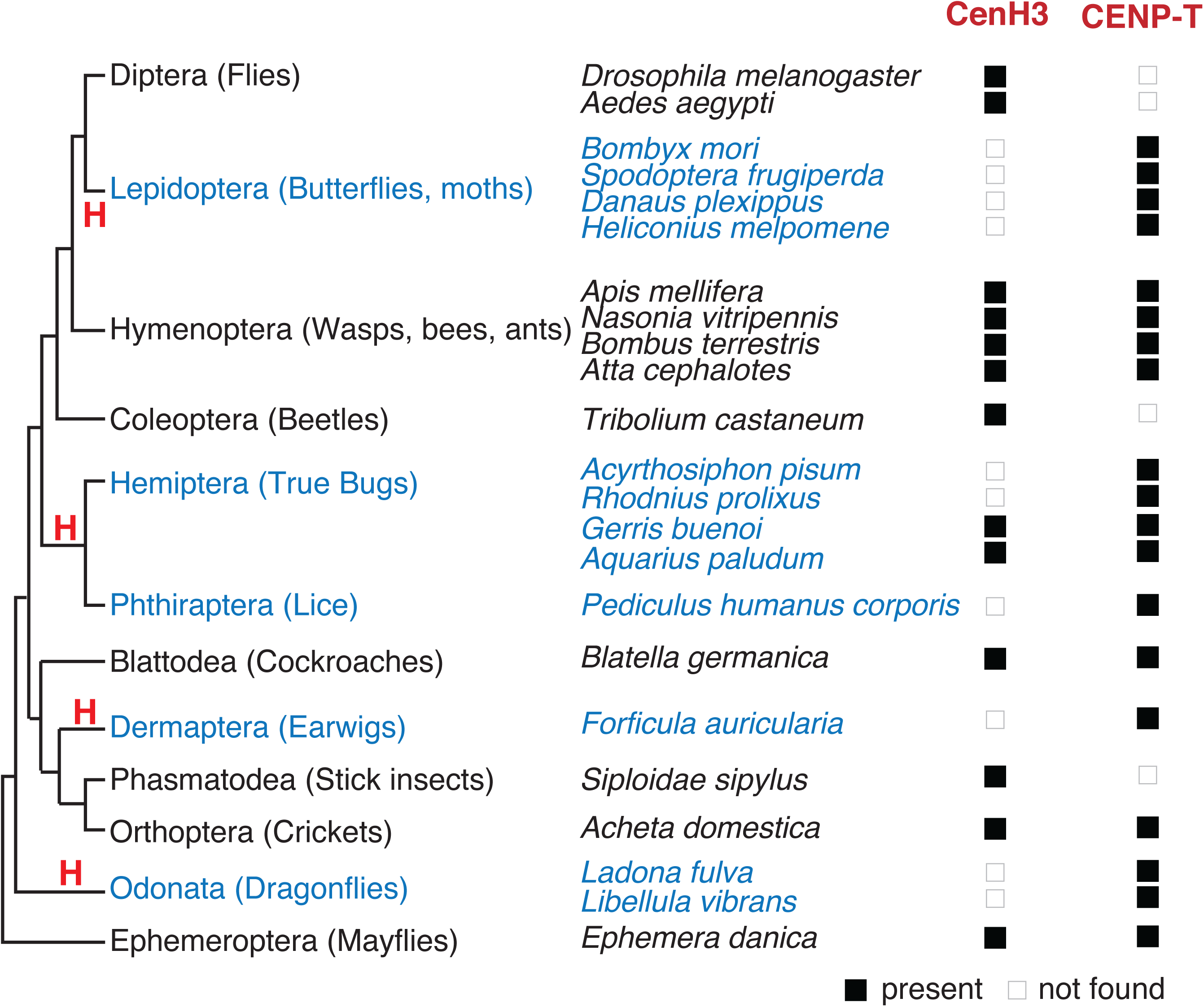
Homologs of CENP-T are present in all other CenH3-deficient insects. Holocentric insect orders and species are indicated in blue, and inferred multiple transitions to holocentric chromosomes are labeled with “H”. Using protein homology searches of genomes or assembled transcriptomes, the ability or inability to find CenH3 and CENP-T homologs are indicated by a black or white box, respectively. Note that while our previous studies did not identify any holocentric insect species that encodes for CenH3 [29], searches in additional organisms revealed the presence of putative CenH3 proteins in water striders (Hemiptera). This result provides new insights into the transition to CenH3-deficient holocentric architectures in that the change in centromeric architecture appears to precede the loss of CenH3, perhaps by providing the necessary conditions to allow the loss of this otherwise essential component.

## Discussion

This study provides new insights into the plasticity of kinetochore formation. CenH3 has long been thought to be the cornerstone of kinetochore formation by mediating the attachment of the kinetochore to chromatin. Its direct DNA binding partner CENP-C in turn contributes to the assembly of other CCAN components and the recruitment of the outer kinetochore in mitosis [5–7,20,21,34]. In addition to these two central components, CENP-T emerged as another core component of the kinetochore capable of bridging chromatin to the outer kinetochore complex in vertebrates and fungi [8,9,24]. Here, the discovery and characterization of CENP-T in CenH3-deficient Lepidoptera provides the first insights into alternative pathways to build a CCAN-based inner kinetochore in a CenH3-independent manner.

Our assays using CENP-T depletion and artificial tethering indicate that the role of the lepidopteran CENP-T in outer kinetochore recruitment is different from that in vertebrates and fungi. In vertebrates, CENP-T contributes to the recruitment of both the Mis12 and the Ndc80 complex by directly interacting with their respective subunits [11,36]. In fungi, the CENP-T N-terminus directly interacts with the Spc24/Spc25 subunit to recruit the Ndc80 complex [8,38]. In contrast, our results show that the lepidopteran CENP-T only contributes to the recruitment of the Mis12 complex. Whether or not CENP-T makes direct protein interactions with any of the Mis12 complex subunits remains to be shown. In addition, other CCAN components also appear to have essential roles in CenH3-independent kinetochore assembly. For example, we find that upon the depletion of CENP-I both the Mis12 and the Ndc80 complex recruitment is abolished, consistent with CENP-I depletion resulting in the strongest mitotic defects. Future studies will aim to dissect the contribution of CENP-I, other CCAN, or even additional kinetochore components with unknown evolutionary relationships (Figure 1) to CenH3-independent kinetochore assembly and chromatin attachment in Lepidoptera.

Considering the essential roles of CenH3 and CENP-C in all other organisms tested, the recurrent loss of these proteins in several insects indicates that the potential for CenH3/CENP-C-independent kinetochore formation might have already arisen in an early insect ancestor. Such potential could be represented in the form of other kinetochore components that evolved the capability to compensate (partially or completely) for CenH3/CENP-C-dependent roles in kinetochore-chromatin attachment and outer kinetochore recruitment. The retention of CENP-T in independently-derived CenH3/CENP-C-deficient insects suggests their important contributions to kinetochore assembly, perhaps by contributing CenH3/CENP-C mediated functions. Notably, given that *D. melanogaster* has lost most CCAN components and solely relies on CenH3 and CENP-C for inner kinetochore assembly, it is unlikely that either protein is dispensable for chromosome segregation in this species.

Our results also exemplify the necessity of experimental data to obtain comprehensive pictures of kinetochore complex composition. The characterization of kinetochore components in additional eukaryotes will enable us to obtain better insights into the sequence divergence of kinetochore homologs. This will allow us to increase the information content of our alignments thereby improving our homology prediction capabilities.

Finally, our studies on kinetochore composition of CenH3-deficient holocentric lepidopteran species, together with the recent finding of a CenH3-deficient monocentric fungus [28,40] that encodes for other CCAN components including CENP-T, show that CCAN-based CenH3 independent kinetochore assemblies are considerably widespread in diverse eukaryotes.

## Acknowledgements

We thank Harmit S. Malik and Steve Henikoff for support on the initial steps of this project, Alexander Schleiffer and Eelco Tromer for helpful discussion on protein homology predictions, Alexander Schleiffer for sharing a collection of representative histone fold proteins, Takahiro Kusakabe for the BmN4-Sid1 line, Tatsuo Fukagawa for the CENP-T-FLAG DT40 line that we used for control experiments, Patricia Le Baccon, Mickaël Garnier and Solène Hervé for help with the microscopic analyses, Ahmed El’Marjou and Carlos Kikuti for help and advice on protein purifications and biochemistry, Carsten Jancke for advice on microtubule stainings, all members of the Fachinetti and Drinnenberg lab for helpful discussions, and Leah Rosin and all member of the Drinnenberg lab for input on the manuscript. NCS receives salary support from Sorbonne Université (2781/2017). This work was supported by “Région Ile-de-France” and Fondation pour la Recherche Médicale grants (to DL). IAD receives salary support from the CNRS. This work is supported by the Labex DEEP ANR-11-LABX-0044 part of the IDEX Idex PSL ANR-10-IDEX-0001-02 PSL, the Institut Curie and the ERC (CENEVO-758757).

## Author Contributions

NCS performed and analyzed the cell biological experiments. JU and EH generated cell lines and performed the IP experiments. TK generated and analyzed the *B. mori* CENP-T knock-out strain and SK supervised the *in vivo* analyses. IAD performed the computational and phylogenetic analyses. GC and IL generated constructs and helped with the biochemical and cell biological experiments. FD carried out the MS experimental work and DL supervised MS and data analysis. IAD supervised the study and secured funding. NCS and IAD wrote the manuscript.

