## Supplemental Material for "CenH3-independent kinetochore assembly in Lepidoptera requires CENP-T"

#### Supplemental Figure Legends

**Figure S1: Kinetochore protein complexes visualized by silver staining.** SDS-PAGE separating protein complexes identified by immunoprecipitation using anti-FLAG affinity steps from Sf9 cell lines stably expressing 3xFLAG-tagged inner kinetochore components (CENP-M, CENP-I, CENP-N and CENP-T) and outer kinetochore components (Dsn1 and Nnf1). Wild-type Sf9 cells (Control) were also used for affinity purifications with anti-FLAG antibodies. Kinetochore proteins used as baits are highlighted in red. The theoretical size of other components identified in the IPs with at least 5 matching peptides are indicated. The identity of several individual bands was determined by mass spectrometry (MS) analysis of digested peptides (blue).

**Figure S2: Prediction of known CENP-T homologs using the C-terminus of the *B. mori* CENP-T protein**. A) and B) Homology searches (arrows) and corresponding *E* values that link the C-terminus of the *B. mori* CENP-T protein to other known CENP-T homologs including *Calypte anna* (Aves), the human CENP-T and the *G. gallus* CENP-T domain structure indicated by species name and UniProt or PDB IDs. While the first jackhammer [51] iteration only picked up insect homologs, the best hits of the second iteration are known CENP-T homologs. The best hit of the HHpred [52] search within the PDB is the *G. gallus* CENP-T. The reciprocal best hit of a search using the *G. gallus* CENP-T HFD helix 3 and extension (amino acid 639-689) against the *B. mori* proteome was the *B. mori* CENP-T. B) Multiple alignment of the C-terminus of vertebrate and insect CENP-T proteins. *B. mori* (above) and *G. gallus* (below) secondary-structure predictions are derived from the Jpred4 (α-helical regions are in red; β-strands in yellow) [75]. Background colouring of the residues is based on the clustalx colouring scheme. Full insect CENP-T sequences and accession numbers if available are listed in Supplementary Table S2.

**Figure S3: Phylogenic distribution of known and insect CENP-T proteins as well as various additional histone fold proteins.** Maximum-likelihood phylogeny of HFD with known CENP-T proteins in orange and newly identified insect CENP-T proteins in red. Bootstrap values above 80 are indicated by blue circles on the branches. The scale bar measures the evolutionary distance in amino acid substitutions per site.

The phylogenetic relationship between the insect proteins (red) and other CENP-T proteins (orange) cannot be unambiguously resolved. Nevertheless, we refer to the insect proteins as CENP-T because of the common sequence architecture with other CENP-T proteins; remote homology predictions identifying known CENP-T proteins as best hits outside of insects (Figure S2) and the experimental evidence of kinetochore participation and outer kinetochore recruitment. Notably, the branch at the root of the insect CENP-T proteins clade indicates a high degree of protein sequence divergence of the insect homologs, which could explain why their putative homology to other CENP-T homologs has been missed in previous searches.

**Figure S4: Absence of detectable lepidopteran CENP-W candidates in *S. frugiperda* FLAG-tagged CENP-T HFD and 2-helix extension (CENP-T-HFDextension-FLAG) immunoprecipitates.** Top: Table lists the number of peptides and coverages of *S. frugiperd*a proteins identified by mass spectrometry enriched in the CENP-T-HFDextension-FLAG IP over the control. The corresponding homologs in *B. mori* and descriptions based on homology predictions are listed alongside. Two small proteins of unknown function are highlighted in green. Bottom: Depletion of *B. mori* homologs of the two small *S. frugiperda* proteins found in CENP-T-HFDextension-FLAG IPs had no detectable mitotic and CENP-T recruitment defect. Graph showing the percentage of mitotic cells (H3S10ph positive cells) three days after RNAi-mediated depletions of corresponding mRNAs (n= number of cells, ± standard error of the mean). Representative images of mitotic BmN4-SID1 cells showing the levels of endogenous CENP-T in cells upon depletion of two small *B. mori* proteins. Scale bar: 10µm.
While we identified two small proteins with sizes similar to known CENP-W homologs in IP experiments pulling down *S. frugiperda* CENP-T C-terminus, RNAi-mediated depletions of *B. mori* homologs do not increase the number of mitotic cells or abolish CENP-T localization to mitotic chromosomes. Both potential protein products (LOC105842400 is annotated as a non-coding RNA) have no detectable similarity to known CENP-W homologs.

**Figure S5: Efficient depletion of kinetochore component encoding mRNAs in BmN4-SID1 lines.** RNA blot analyses showing mRNA levels encoding for various kinetochore components in control (CTRL - RNAi against GFP) and depleted cells (KD). The star indicates the band of the respective mRNAs. A probe against Rpl32 mRNAs were used as loading control. The size marker is in base pairs.
An RNA probe targeting the Dsn1 transcript did not reveal any signal and is therefore not included in this figure. The efficient depletion of Dsn1 protein is confirmed in our IF data analyses (Figure 3). We were unable to assign one band to the CENP-M transcript indicating the possibility of multiple expression isoforms. The signal of all bands was decreased upon RNAi treatment indicating efficient depletion of CENP-M transcript levels.

**Figure S6: Depletion of kinetochore components affects mitotic progression in *B. mori* cells.** A) Quantification of the percentage of mitotic cells (H3S10ph positive cells) five days after RNAi-mediated depletion of various kinetochore components (n= number of cells, ± standard error of the mean). B) Representative images of mitotic cells stained with anti-tubulin used to classify mitotic defects observed at three days after depletion of outer kinetochore proteins Mis12, Nsl1 and Spc24. Scale bar: 10μm.

**Figure S7: Depletion of kinetochore components affects mitotic progression in *B. mori* cells.** A) and B) Zoomed-out versions of representative images of cells stained with anti-tubulin used to classify mitotic defects observed three (A) and five (B) days after depletion of inner and outer kinetochore components. Scale bar: 30μm.

**Figure S8: Depletion of kinetochore components affects mitotic progression in *B. mori* cells.** Representative images of mitotic cells stained with anti-tubulin with mitotic defects observed five days after depletion of inner and outer kinetochore components. Scale bar: 10μm.

**Figure S9: The CENP-T N- and HFD containing C-terminus are both essential for mitotic progression.** A) Schematic representation of the wild-type FLAG-tagged CENP-T (CENP-T-FLAG), FLAG-tagged RNAi-resistant CENP-T (CENP-Tres-FLAG), the N-terminal (ΔN-CENP-Tres-FLAG) and C-terminal truncated RNAi-resistant CENP-T (ΔC-CENP-Tres-FLAG). The recoded region conferring RNAi-resistance to the used dsRNA against CENP-T is indicated in orange (see Methods). B) Representative IF images of mitotic cells expressing CENP-T-FLAG (first row) followed by RNAi-mediated depletion of endogenous and exogenous CENP-T. Representative IF images of cells expressing full-length CENP-Tres-FLAG (second row), ΔN-CENP-Tres-FLAG (third row) or ΔC-CENP-Tres-FLAG (fourth row) RNAi-resistant FLAG-tagged CENP-T followed by RNAi-mediated depletion of endogenous CENP-T. Scale bar: 10μm.

**Figure S10: Validating the specificity of antibodies raised against CENP-T, Dsn1 and Spc24/25 complex.** A) and B) Top: Schematics showing protein fragments used for antibody generation against lepidopteran CENP-T and Dsn1 homologs. Bottom: Mass spectrometry results identifying proteins (> 5 peptides) present in immunoprecipitates from *B. mori* soluble extracts using the antibody raised against the CENP-T N-terminus or Dsn1, respectively, but not in immunoprecipitates using the Sigma M2 antibody as a control. C) Top: Schematics showing protein fragments used for antibody generation against the *B. mori* Spc24/25 complex. Bottom: Commassi staining and western blot analyses of different IP fractions pulling down *B. mori* His-Spc24/Spc25 protein fragments expressed in Sf9 cells over Nickel-NTA columns. Two bands of the expected size for the His-Spc24 and Spc25 fragments are labelled with a star on the commassi-stained gel. Increased concentrations of immidazol are indicated in mM for the wash and each elution step. The α-His tag immunosignal confirms the presence of the His-Spc24 fragment in several elution fractions. A band of the same size in the same elution fractions is also recognized by the custom-made antibody against *B. mori* Spc24/Spc25 fragments. An additional, much fainter, band corresponding to the size of the *B. mori* Spc25 fragment is also detected. D) Immunofluorescence signals for CENP-T, Dsn1 and Spc24/25 in combination with histone 3 serine 10 phosphorylation (H3S10p) in mitotic BmN4 cells with and without RNAi-treatment against the corresponding mRNAs. Images from this panel are the same as in Figure 3. White bar represents the scale of 10 µm.

**Figure S11: Depletion of additional outer kinetochore components abolishes recruitment of their respective complex members in *B. mori* mitotic cells.** A) Representative images of mitotic BmN4-SID1 cells showing the levels of endogenous CENP-T, Dsn1 and Spc24/25 with and without depletion of outer kinetochore components (Mis12, Nsl1 and Spc25). Scale bar: 10µm. B) Quantification of mean fluorescence intensity for anti-CENP-T, anti-Dsn1 and anti-Spc24/25 signals upon kinetochore depletion compared to the control. Statistical significance was tested using the Mann and Whitney test ($p < 0.0001$ for four stars, $0.001 < p < 0.01$ for two stars).

#### Methods

##### Alignments and phylogenetic analyses

For Figure 1C and Figure S2C, sequences were aligned with MAFFT on the EMBL-EBI web interface [35], and visualized and processed with Jalview [37] (Clustalx colouring scheme). Secondary structures were predicted using Jpred4 [39]. For Figure S3, histone fold domains were aligned using MAFFT [45,47]. A maximum likelihood phylogeny was build using PhyML 3.0 [41] with automatic model selection [42], default parameters and 100 bootstrap permutations. The tree was visualised using iTOL vs4 [43]. Sequences used for the phylogeny are listed in DatasetS1.

Primary sequence analyses of the *B. mori* and *G. gallus* CENP-T (NP_001263242.1) for Figure 1C were performed using fLPS [44] with default parameters. The Arginine-rich N-terminal regions are located between amino acid 4 and 32 of the *G. gallus* CENP-T (*E* value $6.0x10^{-7}$); and 20 and 41 of the *B. mori* CENP-T (*E* value $4.7x10^{-5}$). The proline-rich regions are located between amino acid 493 and 521 of the *G. gallus* CENP-T (*E* value $3.4x10^{-8}$); and 850 and 899 of the *B. mori* CENP-T (*E* value $3.7x10^{-4}$).

##### Homology predictions

The annotation of the *S. frugiperda* “corn strain” were used for all computational analyses [46]. In case annotations were missing or wrong we complemented the data using EST data [48] or our own analyses. Homologs of *S. frugiperda* proteins identified in the kinetochore IPs were predicted in *B. mori* proteome [49]. Both *S. frugiperda* and *B. mori* proteins were used for homology searches. BLASTP and PSI-BLAST searches were performed in the NCBI non-redundant protein database [50]. Jackhmmer searches were performed on the HMMER webserver [51] against reference proteomes as current as September 2019. HHpred version 3.2.0 searches were performed against the PDB_mmCIF70_3_Aug [52]. Coiled-Coil domains were predicted using PCOILS (window 28) [59,61,52].

*CENP-T*

For iterative HMM profile searches only the C-termini containing the HFD and 2 helix extension of the *B. mori* or *S. frugiperda* proteins were used.

>B_mori_CENPT_CTERM

```
TTKRLYKYLEDKLEPKYDYKARVRAEKLVETIYHFTKEVKKHEVAPNDAVDVLKHEMARLDIVKTHFDFYQFFHD
FMPREIRVKVVPDIVNKITIPRNGVFSEILSGHAVHA
```

>S_frugiperda_CENPT_CTERM

```
ITKRLYKFLETKLEPKYDYKARVRAEKLVETIYHFAKDLRRHDVAPTDAVDVLKHELARLEVVQTHFEFYEFFHE
FMPREVRVKVVPDIVNKIPLPRHGVFSDILRGNNVQG
```

HHpred searches using the C-terminus of the *B. mori* and S*. frugiperda* CENP-T protein identify the C-terminus of the *G. gallus* CENP-T with high probability 94.56 % and 94.2 % respectively (Figure S2). HHpred searches using the full-length protein also identifies the *G. gallus* CENP-T as a best hit though with lower probabilities (51.9 % for the *B. mori* and 50% for the *S. frugiperda* protein). In this search, the alignment between the lepidopteran protein and *G. gallus* CENP-T is restricted to their C-termini. A reciprocal HHpred search using the *G. gallus* HFD helix 3 and CENP-T extension identified *B. mori* protein as best hit (Figure S2).

Known vertebrate CENP-T proteins can also be identified as best hits after 2 jackhmmer iterations using the *B. mori* CENP-T C-terminus (Figure S2). Additional arthropod CENP-T from the whiteleg shrimp (*Penaeus vannamei*) and the pill woodlouse (*Armadillidium vulgare*) could be predicted after 4 jackhmmer iterations. Finally, phyre2 predictions [53] using the full-length *B. mori* CENP-T protein predicts the known structure of the *G. gallus* CENP-T C-terminus with high confidence (91.5%) while low sequence identity of the aligned regions (14%).

Iterative blastp and tblastn searches using lepidopteran CENP-T proteins against select annotated insect proteomes, genome assemblies and assembled transcriptomes revealed additional orthologs of CENP-T proteins in insects (Table S2).

*KWMTBOMO14835 and GSSPFG00001205001*

BLASTP searches using both proteins against the nr database revealed only hits in Lepidoptera. Iterative PSI-BLAST searches and jackhmmer searches using both proteins also revealed no homologous hits in other insect orders. HHpred searches of GSSPFG00001205001 did not reveal any hits with high probability. However, HHpred searches identified similarity between the N-terminus of KWMTBOMO14835 and resolved structure of the *S. cerevisiae* Ctf19 protein (CENP-P) [54] as best hit with high probability (93.58%) while only 13% amino acid identity. The aligned region also includes parts of the tandem RWD domains from *S. cerevisiae* Ctf19. The N-terminus of KWMTBOMO14835 has predicted coil-coiled regions (PCOILS, window 28). HHpred predictions of the trimmed protein without the coiled-coil region reduced the probability of similarity to Ctf19 to 43.96%, which was also no longer the best hit.

We therefore refrain from assigning homology to Ctf19 without further functional or structural validations and refer to the lepidopteran proteins as coiled-coil RWD-like proteins.

*KWMTBOMO09290 and GSSPFG00019785001*

Both proteins contain N-terminal coiled-coil domains (PCOILS, window 28). HHpred searches of both proteins without the N-terminal coiled-coil regions revealed similarities to RWD domains of several kinetochore components including the resolved structure of the *G. gallus* Spc24 globular domain [9] and Mcm21 (CENP-O) subunit of the budding yeast Ctf19 complex [55,54] with high and similar probabilities. Iterative jackhmmer searches of the trimmed *B. mori* protein without the N-terminal coiled-coil region identified hits in Hymenoptera in the second iteration including *Camponotus floridanus* (E2AEP7, *E* value 0.0017). The third iteration identified additional hits in the blattodean *Zootermopsis nevadensis* (A0A067RMY5, *E* value 0.00028), the arthropod *Daphnia magna* (A0A0P5Q3Q5, *E* value $6.5 \times 10^{-7}$) and predicted known vertebrate CENP-O proteins in *Anabis testudineus* (A0A3Q1H0S6, *E* value 0.0023 and A0A3Q1H0T5, *E* value 0.0037). The fourth iteration identified the human CENP-O (*E* value $1.6 \times 10^{-13}$).

Given the similarity to multiple RWD containing kinetochore proteins and the lack of experimental evidence we only refer to the lepidopteran proteins as coiled-coil RWD-like proteins.

*KWMTBOMO06154 and GSSPFG00011797001*

HHpred searches did not identify high scoring hits for neither one of the two proteins. GSSPFG00011797001 identified the recently resolved structure of the fungus *Thielavia terrestris* CENP-H [56] but not as the best hit and with only 36 % probability. Still, given the presence of CENP-I, CENP-M and the putative CENP-K, it will be worthwhile to test if these proteins are part of a lepidopteran CENP-H-I-K-M complex. Hits in other insect orders could not be predicted using psi-blast or jackhmmer.

*LOC101741561 and GSSPFG00006732001*

The *B. mori* homolog is not part of the most recent annotations [49] , we therefore refer to the ID of previous annotations present on NCBI Genebank in Figure 1.

GSSPFG00006732001 is likely misannotated because most other lepidopteran homologous proteins including LOC101741561 only align to the C-terminal part of the protein. We derived a new consensus sequence by aligning available EST data [48]. The encoded ORF translates into the following protein:

>*S. frugiperda* CENP-K

```
MSSDRNKETRDAVKREIKEIQARCKHEWNIIDNSPLDAPNIDLEQINEKALCYIEGLGTGIQNSNTPITADDNLL
TSQFLKEIRDKTGQVEEYTAFVRGSIHDIDAEINRLQTLIKITQEAKARPMLNKCEVQPEHVHRAKERFQVMKNE
LHSLIHSLFPNCDSLIIETMGQLMAEHLNEESNGYIPVTAETFQIIELLKDMKIVTVNPYNKLEVKLSY
```

One of the EST that contain the full ORF is Sf2H05447-5-1 that is listed in Figure 1B.

Iterative PSI-BLAST searches using the *B. mori* protein revealed hits in Hymenoptera including *Athalia rosae* (LOC105689000, *E* value $5x10^{-5}$) after the 1st iteration. The 2nd iteration revealed hits in Hemiptera including *Bemisia tabaci* (LOC109036758, *E* value 0.003), the Blattodean *Zootermopsis nevadensis* (LOC110834781, E value $2x10^{-11}$) and the mollusk *Mizuhopecten yessoensis* (centromere protein K-like, *E* value 0.001). The human CENP-K was identified after the 3rd iteration. These analyses are consistent with previous predictions identifying CENP-K in *B. mori* [30].

Jackhmmer searches of a N-terminal truncated version of the *B. mori* protein without a coiled-coil region also revealed a hit in the phthirapteran *Pediculus humanus corporis* after the 3rd iteration (E0VW71, *E* value $4.1x10^{-9}$).

*KWMTBOMO11351 and GSSPFG00010096001*

These are orthologs of *Drosophila melanogaster* bellwether, the alpha subunit F0F1 ATP synthase subunit alpha recently been shown to interact with Cid during *Drosophila melanogaster* male meiosis [31]. Orthologs are present across insects and diverse eukaryotes.

*KWMTBOMO11557 and GSSPFG00019290001*

HHpred searches aligns both proteins to the structure of the CHRAC 14 HFD [57] as best hits but low probabilities (Probability 49.3 % for KWMTBOMO11557 and 39.1% for GSSPFG00019290001). Hits in other insect orders could not be predicted using psi-blast. Jackhammer searches of both proteins converged after 2 iterations.

*Sf2H06980-5-1 and KWMTBOMO014775*

The *S. frugiperda* protein is not part of the *S. frugiperda* corn strain annotations. We therefore provide the ID of one of several ESTs which contain the ORF [48]. Both *B. mori* and *S. frugiperda* proteins contain C-terminal coil-coiled regions. HHpred searches using only the N-terminal first 65 amino acids identifies the known structure of the human Nsl1 [58].Though the human Nsl1 was not the best hit for neither one of the two lepidopteran proteins, given the abundance of the protein in the Nnf1 and Dsn1 IPs we refer to it as Nsl1 candidate. These data suggest that as in other eukaryotes, the lepidopteran Mis12 complex contains four instead of only three components as recently described [33].

*Hemipteran CenH3*

Hemipteran CenH3 fragments in *Gerris buenoi* genome assembly [60] and *Aquarius paludum* assembly (A. Khila, unpublished data) could be identified in TBLASTN searches using *D. melanogaster* H3 as queries.

>Aquarius_paludum_Embryo_Assembly50_c84768)g2_il
RRRSGVVALREIRHLQKSTNLLIPKLPFMRIVQEILQSYSTEPYRIQSRALEALQEMTEILMVDLFSEAILCCIH
AKRKTIMVQDMRLARRIRG

>Gerris_buenoi_ KZ651887.1
RRRSGVVALREIRHLQKSTNLLIPKLPFMRIVKEILQSYSTEPYRLQTQALEALQEMTEILMVDLFSEAILCCLH
AKRKTIMVQDMRLARRIRG

**Plasmid construction**

A list of plasmids generated in this study is provided in Table S5. For the kinetochore IPs, full-length ORFs of *S. frugiperda* CENP-M, CENP-N, CENP-I, CENP-T, Dsn1 and Nnf1 fused to 3xFLAG tags were cloned into pIBV5 (12550018, Invitrogen) using the Gateway system (11791019, Invitrogen) from pENTR vectors (K240020, Invitrogen). The genes encoding for *S. frugiperda* CENP-T, CENP-I and CENP-N are not correctly annotated in the current assembly [46] as inferred by aligning the protein products to the *B. mori* homologs. We inferred the correct ORF sequences from EST data or sequencing the amplified PCR fragments from Sf9 cDNA.

The following are the correct ORF sequences that were used for all experimental analyses:

>*S. frugiperda* CENP-I
MADVDEIIDYIKSLKKGFDKDLFQNKIDELAYAVDTTGILYNDFHTLFKVWLNLSIPITKWVSLGACLVPQNIVEDRTVEYALSWMLSNYEDQS
TFSRIGFLLDWLTAAMECECIEMETLDMGYDVFYVSLTYETLTPHAVKLVYTLTKPVDVTRRRVLELLDYARKREAKKNMFRQLXVLLGLFKSY
KPECVPEDIPAISIHAAFKKINPDLLARFKRNQENRNSVRRERHHLTWINPINSDRGRNKKIDPLVPNVEFLNIGSKQYAEKEHQKNFLDFTDP
VSVLQCSVQRSTSRPARIRALLCNVTGVALLAVASHTEQEFLSHDLHHLLNSCFLNISPHSYREKQDLLHRLAVLQHTLMQGIPVITRFLAQYL
PLWNERDYFAEILELVQWVSLDSPDHVTCVLEPLARAYHRAQPIEQCAILRSLNHMYCNLVYASTRKRHHFMGTPPSPQVYALVLPKVATAISD
MCDKELQVNPEQMVVLHSGVQGAAARARGEARGGAAAGALAPRLALALPLLASSAALLDSVAELMILYKKIFTTAKQTNVITNTDTFEKQMQVL
EAYTSDLINCLYSEGALSDRNLGFVFSKLHPQLVEKLGSLMPDVDAKLSIRNSIAFAPYTYIQLDAIDHRDADNKLWFNAVIEQEFTNLSRFLK
RAVTELRYQ

>*S. frugiperda* CENP-T
MPKTKIPSPARPQGATTPKRKKSRGSRSVCSPALSTASGRLTSLIDQFKEKNMESFALSPLNRRSILNDDTAIEEPRRQSWWKKLKEDSHEIME
VLEENKVADSGNNAIEELIDIEVLSQEKKEYTLDLPESSDNESINSIVLPQRKLFTQKENKPQKKFGQFSDNRETLAKLHKTNTQGDKTVNVGT
RELFQNAKRSKPIFPAALLNISPNKTAMDKTKENILPEPKVRNIFGNRPANKRKNMFADFVVSESEDEIPELQPRVFGFQKKLEQRRISSISKG

REGSPASSITTDMEMDDWKLLPSSTMVENQLEDIVAGHTPKRARLSKLSEAKESEAGTLQTNTTGDKTKSSNKSIMSKNKSKSSPDAKDKSLNS
SRTTRSMLKNASKLEENTEPKQDTKTQSKISTKDNALDVSLSKATTKQNTSLNKSLRLNKSRTRNSSKLNEEEMETDENVVKNAINDATRVISK
NASSAAVASKSINEKERTITLKVHDKASETGSNKEEDDNNFVLQYEDEIVEEATDHNQINENKTTKIKNRNTIVIENDQETDVNESKSNKSIQK
EPKQAKTVEESVESQNEQSGNDNIDGNEIEENAIEQNHESHDGEKISRELNEESNNSEGNDERNVTKDNKDQENDEEINESQENDEEINLSQES
NRDEAVLSRVDEENESEVNEESENEDDVNESNESEEQNESQEVEAEVDESEEIEEQNESQEIENEANESQEVNEEADDSKVEENASEDEEENQE
VENDEEENAEESQEVDNEESQEIENQEEFEVSQESEHEVSQEVENEEENEEENDEGDQEVEQEEENQSEDDVEDQVEDQSQEIENDEENDEEEN
EESQEIENDEENVVESEAEVNESQEIEGDQEMDDSDEADRAIASDPEISDQDDNGADEMEVDDDDDNNEQESENEEETEGNQEESQEEQQEPSA
EEIEESPNVTHDTTGRHRQKVLKSPEAILHDKTNQMDSFTAKGRNTSIRKTKSMIKNLNIRPSLAPQRDSLAFSDGTRDSSAEGSGWDSHRTTR
KTLRQTFGKDFTPRKSLRALVMEKSAKRQTEHMDLHSETSKYPQANSTELAEDSNGHVDDFVESDHEVSRRTRQTTLETYLQKIKQKNLENKVK
MEELVRNSLKAPARDTLSLFKVPNKPAPRRLKPTQNKPRTQVKAITAFGELPTEVIEDMKYKPPKRFQPTNASWITKRLYKFLETKLEPKYDYK
ARVRAEKLVETIYHFAKDLRRHDVAPTDAVDVLKHELARLEVVQTHFEFYEFFHEFMPREVRVKVVPDIVNKIPLPRHGVFSDILRGNNVQG

>*S. frugiperda* CENP-N

MPLEVFCVTALPEFGVRWRELSKAVRNARLPMQPASVARVLCRHVSKALTADEIEDIVARLRLKLVATQPRTWHVIRLSEKTTEEPVTLTMRAV
PGRITQALRKSKKTMRPEVQTVLLGDMLYLSIQLVSEHKSGSALYVATPPGEPVALVSSVNMVGLIKATVEGLGYKSYEIADLHGRDIPSLLRI
NDRAWNTNADHLAEIPEYAPTPIITETGIDYTYKAYDENYVENILGPNPPKITDLTIKTSKSFFDRSRLDKNINITINIKTEDLAKSLKCWVSK
GAIAPTSDLIKIFHQIKSNKISYTREDD

To express the C-terminus of *S. frugiperda* CENP-T a 166 amino acid C-terminal fragment that contains the HFD and CENP-T extension was isolated to generate the *S. frugiperda* CENP-T-HFDextension-FLAG pIBV5 expression clone.

For LacO/LacI tethering assays, pVS1-LacO (33143, Addgene) was modified to insert the blasticidine resistance cassette from pIBV5 into the BamH1 and Sac1 sites. The construct pVS1-LacO-Blasticidin was transfected into Sf9 cells to establish a stable line. To generate LacI-GFP pIBV5, the LacI-eGFP insert was amplified from eGFP_N1_LacI pDEST (kind gift from the Almouzni lab, Nuclear Dynamics unit, UMR3664, Institut Curie, Paris, France) and cloned into pENTR and pIBV5. To generate *S. frugiperda* CENP-T-GFP-LacI pIZV5, the full-length coding sequence of *S. frugiperda* CENP-T from *S. frugiperda* CENP-T-FLAG pIBV5, lacI from pCMV-lacI (kind gift from the Almouzni lab) and eGFP from eGFP_N1_LacI pDEST were isolated by PCR and assembled into the shuttle vector pRS416 (kind gift from Heloise Muller, Nuclear Dynamics unit, UMR3664, Institut Curie, Paris, France) using the yeast-based homologous recombination-based DNA assembly, described in detail in elsewhere [62,63]. The *S. frugiperda* CENP-T-GFP-LacI was isolated from this plasmid and cloned into pIZV5 (V800001, Invitrogen) using KpnI and XbaI. To generate the *S. frugiperda* ΔN-CENP-T(216-1314)-GFP-LacI pIZV5, a truncated *S. frugiperda* CENP-T-GFP-LacI fragment (without the part coding for the first 215 amino acids, which were used as epitopes for the lepidopteran CENP-T antibody generation) was isolated PCR and cloned into pIZV5 using BamHI and XhoI sites.

To generate the *B. mori* CENP-T-FLAG pIZV5 construct, the coding sequence of *B. mori* CENP-T was isolated from a BmN4 cDNA library, fused to the 3xFLAG tag by PCR amplification and cloned into pIZV5 using BamHI and XhoI. To generate the *B. mori* CENP-T RNAi-resistant clone (*B. mori* CENP-Tres-FLAG pIZV5), we ordered a Gblocks gene fragment (IDT) to recode an internal sequence of *B. mori* CENP-T from amino acid 221 – 334. A 5' fragment of *B. mori* CENP-T that contains the recoded part was then assembled with the 3' fragment of *B. mori* CENP-T fused to a 3xFLAG into pRS416 using yeast-based homologous recombination and subsequently cloned into the into the BamHI and XhoI sites of pIZV5. To generate the *B. mori* ΔN-CENP-Tres-FLAG pIZV5 and *B. mori* ΔC-CENP-Tres-FLAG pIZV5, *B. mori*

CENP-Tres-FLAG pIZV5 was used as a template to isolate 5' or 3' truncated fragments to clone those into pIZV5 using BamHI and XhoI. The constructs expressed CENP-T without the first 200 or last 112 amino acids, respectively.

To generate protein antigens for antibody production, we purified protein fragment expressed in bacteria. To generate the *S. frugiperda* SUMO-6xHis-CENP-T (1-122) pT7 and *B. mori* SUMO-6xHis-CENP-T (1-218) pT7 bacterial expression constructs the N-terminal domains of *Spodoptera frugiperda* (1-222aa) and *Bombyx mori* (1-218aa) CENP-T were cloned into the pT7-His-SUMO vector (gift from Ahmed El-Marjou, recombinant protein platform, Institut Curie) which included N-terminal 6X His and SUMO tags using Gibson Assembly.

To generate the *B. mori* 6xHis-Spc24(73-162)-Spc25(70-211) pRSF-DUET1 bacterial expression constructs, the globular domain of *B. mori* Spc24 (73-162aa) was cloned with an N-terminal 6X His-tag into the first cassette of the pRSF-Duet1 vector using the Bam HI and PstI restriction sites. For dual expression of Spc24 and Spc25, the globular domain of *B. mori* Spc25 (70-211aa) was also cloned into the second cassette of the same pRSF-Duet1 vector using the Nde I and Xho I restriction sites. To generate the *B. mori* 6xHis-Spc24(73-162)-Spc25(70-211) pFASTBac-Dual for the recombinant baculovirus generation to evaluate the Spc24/25 antibody, the Spc24 and Spc25 fragments were cloned into the BamHI and PstI, and KpnI and XhoI sites of pFASTBac-Dual, respectively.

To generate the *B. mori* 6xHis Dsn1(1-100) pRSF-DUET1 and *S. frugiperda* 6xHis Dsn1(1-99) pRSF-DUET1, N-terminal region of *B. mori* (1-100aa) and *S. frugiperda* (1-99aa) Dsn1 cloning into pRSF-Duet1 with N-terminal 6X His-tag BamHI + PstI.

**Cell culture conditions**

Cultured silkworm ovary-derived BmN4 (ATCC CRL-8910) and BmN4-SID1 cell lines [64] were maintained in Sf-900 II SFM medium (10902-088, Gibco) supplemented with 10% fetal bovine serum (CVFSVF0001, Eurobio), antibiotic-antimycotic (15240-062, Gibco) and L-glutamine (25030-024, Gibco) at 27 °C. Sf9 cells (12659017, Gibco) were maintained in Sf-900 II SFM medium (10902-088, Gibco) supplemented with antibiotic-antimycotic (15240-062, Gibco) and L-glutamine (25030-024, Gibco) at 27 °C.

**Construction of stable cell lines**

Around 1-5 μg of plasmid DNA was transfected into $10^6$ BmN4 or Sf-9 cells using Cellfectin II (10362100, Gibco) according to the manufactor's instructions. For IF experiments cells were grown on coverslips before transfections. For the generation of stable cell lines, antibiotics were added 48 hours after transfection (300 μg/ml Zeocin (R25001, Gibco) or 40 μg/ml Blasticidin (R21001, Gibco). Selection was continued until no viable untransfected cells were observed.

**Protein expression and affinity purification**

The *E. coli* Bl21DE30 plys pRare (Merck) was transformed with bacterial expression vectors containing the recombinant genes. Bacterial cultures (4 L) were initially grown at 37 °C, 220 rpm and then induced with 0.5 mM IPTG (EU0008-A, Euromedex). Following expression at 37 °C for 4 h, cultures were centrifuged at 6000 *xg* for 15 min to pellet the cells. Cell pellets were resuspended in Wash Buffer (20 mM Tris pH 8, 300 mM NaCl, 10 mM $MgCl_2$, 25 mM imidazole). The resuspended pellets were incubated at 4 °C on a roller with the addition of one tablet cOmplete Protease Inhibitor Cocktail (11697498001, Roche), Triton-X-100 (1% final concentration), and lysozyme (1 mg/mL final concentration). The mixtures were sonicated at 30% amplitude for 8 min total (30 sec pulses) in a Branson Digital Sonifier SFX550 (Branson Ultrasonics Corp.). Cell lysates were centrifuged at 30 000 *xg* for 1 h at 4 °C. The soluble fraction was removed and passed through 0.45 µm filter units. The lysates were loaded onto Protino Ni-TED 1000 columns (Machery-Nagel) that were pre-equilibrated with Wash Buffer. The columns were washed with 2-3 column volumes of Wash Buffer and proteins were eluted with the following buffer (20 mM Tris, 250 mM imidazole, 300 mM NaCl, pH 8.0). The SUMO tags were cleaved from the CENP-T proteins by digesting the eluates at 4 °C with SUMO-Protease (enzyme produced by Ahmed El Marjou, recombinant protein facility, Institut Curie) (final concentration 0.6 µg/mL). The eluates were loaded and migrated in Bolt 4-12% Bis-Tris Plus denaturing gels (Invitrogen). The correct-sized bands corresponding to the proteins without the SUMO tag were excised from the gels and sent to Covalab (Villeurbanne, FR) for generation of antibodies in rabbits.

**Validation of lepidopteran CENP-T and Dsn1 antibody specificity by IP and MS**

For each IP experiment, one confluent flask of BmN4 cells was used. Immunoprecipitation was performed as described previously [65] omitting the cross-linking step and with some modifications. Chromatin was digested using I UNIT of MNase (N3755-500UN, Sigma) followed by mild shearing and solubilization step using the Covaris E220 Evolution ultrasonicator (150 sec, peak power 75, duty factor 10, cycles/burst 200). The soluble extract was added to 50 µl magnetic Dynabeads Protein A (10002D, Invitrogen) covalently conjugated to 5 µl of rabbit polyclonal anti-CENP-T or anti-Dsn1 immunoserum or 10 µg ANTI-FLAG M2 antibody (F1804, Sigma) as a control. Immunoprecipitation was performed for 15 minutes at room temperature and samples were washed three times in IP dilution buffer (1% Triton X-100, 2 mM EDTA, 150 mM NaCl, 20 mM Tris-HCL (pH 8.1)). Samples were digested on beads for MS analyses (Figure S10A and B) (see below).

**Validation of *B. mori* Spc24/25 antibody specificity**

The 6xHis-Spc24(73-162)-Spc25(70-211) pFASTBac-Dual construct was used to generate recombinant baculovirus DNA using the flashBAC technology, flashBACULTRA™. Recombinant baculoviruses were amplified in Sf9 cells to generate high-titer virus stocks. 500 µl of the recombinant Spc24/Spc25 expressing baculovirus stock (MOI 100) were used to infect 50ml Sf9 cells ($10^6$ cells/ml) for 3 days. To obtain total cell extracts, the cell pellet was resuspended in buffer A (20 mM Tris pH 8, 300 mM NaCl, 5% glycerol, one tablet cOmplete Protease Inhibitor Cocktail (11697498001, Roche) and 20µl DNase I (04716728001, Roche)) and incubated for 20 min at 4°C. The samples were sonicated at 20% amplitude for 3 min and 30 sec total (30 sec pulses) in a Branson Digital Sonifier SFX550 (Branson Ultrasonics Corp.) and centrifugated for 30min at 8000 rpm, at 4°C. The supernatant was used for the subsequent experiments. The lysates were loaded onto Protino Ni-TED 1000 columns (Machery-Nagel) that were pre-equilibrated with Wash Buffer. After a 30 min incubation at 4°C, the columns were washed in Wash Buffer with 20 mM imidazole followed by several elution steps with increasing concentrations of imidazole (50 mM, 100 mM, 200 mM and 500 mM). Proteins were separated on 4-20% Tris glycine gels (XP04200BOX, Invitrogen) and visualized using Instant*Blue*™ (ISB1L, Sigma).
For protein blot analysis, samples were separated on Bolt 4-12% Bis-Tris Plus gels (NW04120BOX, Invitrogen) and transferred to a PVDF membrane (170-4272, Bio-Rad) using the Trans-Blot Turbo Transfer System (1.3 A, 25 V, 10min). The membrane was blocked using the Odyssey Blocking buffer (927-50000, LI-COR) before primary antibody incubation (rabbit polyclonal anti-Spc24/Spc25, 1:1000 dilution and mouse monoclonal anti-6xHis, 1:1000 dilution (ab18184, Sigma)) and secondary antibody incubation (IRDye 680RD Goat anti-Rabbit IgG (926-68071, LI-COR), IRDye 800CW donkey anti-mouse IgG (926-32212, LI-COR), dilution 1:10000). The signals were visualized on an Odyssey LI-COR scanner (Figure S10C).

**Affinity co-immunoprecipitations**

Cultures of the following Sf9 strains expressing full length or partial *S. frugiperda* kinetochore proteins fused to a C-terminal or N-terminal 3XFLAG tags or control wild-type Sf9 cells were grown to exponential phase in Sf900 II Media (Gibco): CENP-I, CENP-M, CENP-N, CENP-T, CENP-T-HFDextension, Dsn I and Nnf1. For each strain, 3 x $10^9$ cells were harvested by centrifuging for 10 min at 300 *xg*, and the pellets were washed twice in cold PBS. The pellets were resuspended in 5 mL HDG150 Buffer (20 mM HEPES pH 7.0, 150 mM KCl, 10% glycerol, 0.5 mM DTT, 1 tablet cOmplete Protease Inhibitor), and then cells were disrupted with 50 strokes in a dounce homogenizer at 4 °C. The dounced fraction was centrifuged at 1700 *xg* for 10 min at 4 °C, and the nuclei (lower fraction) were gently resuspended with 5 mL HDG150 Buffer and re-centrifuged in the same conditions. The nuclei were resuspended in a final volume of 5 mL using HDG150 Buffer. Nuclear extracts were prepared by passing the nuclear fraction

10 times through a 20G 1 ½" needle (0.9 x 38 mm) and then 5 times through a 25G 3/8" needle (0.5 x 16 mm). The nuclear extracts were centrifuged at 20 000 *xg* for 10 min at 4 °C in microcentrifuge tubes. The pellets were resuspended in HDG150 Buffer and pooled together at a final volume of 5 mL. To prepare the chromatin, the nuclear extracts were digested with ~40 units MNase (N3755-500UN, Sigma) for 1 hour at 4 °C on a roller, 3 mM $CaCl_2$ was added to the digestions. The MNase digestions were stopped by adding 250 µL of 0.2 M EGTA. To solubilize the digested chromatin, 10 mL of HDG400 Buffer (20 mM HEPES pH 7.0, 400 mM KCl, 10% glycerol, 1 mM DTT, 0.05% NP-40, 1 tablet cOmplete Protease Inhibitor) was added to the samples and incubated for 2 h at 4 °C on a roller. The samples were centrifuged at 8000 *xg* for 10 min at 4 °C. The supernatants were saved to bind to the Anti-3XFLAG M2 beads (M8823, Sigma). The M2 magnetic beads were prepared according to the manufacturer's recommendations. The digested chromatin samples were incubated with 150 µL Anti-3XFLAG M2 beads. The beads were washed four times with 1 mL HDGN320 Buffer (20 mM HEPES pH 7.0, 320 mM KCl, 10% glycerol, 1 mM DTT, 0.05% NP-40, 1 tablet cOmplete Protease Inhibitor). For proteomic analyses of full-length kinetochore protein IPs, beads were boiled in sample buffer to first run those into gels (see below). For proteomic analyses of the CENP-T-HFDextension IP, proteins were directly digested on beads (see below). For the silver stainings of Sf9 kinetochore IP samples, M2 beads were incubated with FLAG peptide (F4799, Sigma) diluted to a final concentration of 150 µg/mL in 750 µL TBS Buffer (50 mM Tris-HCl, pH 7.4, with 150 mM NaCl) for 1 h at 4 °C on a roller. The supernatants were removed and the eluates were concentrated in Amicon Ultra-0.5 mL 3K MWCO filters (Sigma) according to the manufacturer's recommendations. The other set was loaded and migrated on Novex 16% Tris Glycine Precast Gels (Invitrogen). Silver stains were performed on the gels using the Pierce Silver Stain Kit (24612, Thermo Fisher Scientific) according to the manufacturer's instructions. Several bands were excised for mass spectrometry.

**Proteomics and Mass Spectrometry Analysis.**

IP enriched proteins of full-length kinetochore protein IPs and control samples were separated on 10% SDS–PAGE gels (Invitrogen) and stained with colloidal blue (786-35, LabSafe GEL BlueTM GBiosciences). SDS-PAGE was used with short separation as a clean-up step, and 4 gel slices were excised. Gel slices were washed and proteins reduced with 10 mM DTT prior to alkylation with 55 mM iodoacetamide. After washing and shrinking the gel pieces with 100% MeCN, in-gel digestion was performed using trypsin/Lys-C (V5071, Promega) overnight in 25 mM $NH_4HCO_3$ at 30 °C. Peptides were then extracted using 60/35/5 MeCN/$H_2O$/HCOOH and vacuum concentrated to dryness. Protein on beads samples (CENP-T-HFDextension IP and CENP-T and Dsn1 antibody validations) were washed twice with 100 µL of 25 mM NH4HCO3 and submitted to on-beads digestion with 0.2 µg of trypsin/Lyc-C for 1h. Digested sample were then loaded onto homemade C18 StageTips for desalting and peptides were eluted using 40/60 MeCN/H2O + 0.1% formic acid and vacuum concentrated to dryness. Gel samples were chromatographically separated using an RSLCnano system (Ultimate 3000, Thermo Scientific) coupled to an Orbitrap Fusion mass spectrometer (Q-OT-qIT, Thermo Fisher Scientific with a Nanospay Flex ion source (Thermo Scientific) and bead samples were also analyzed with a Q Exactive HF-X mass spectrometer. Peptides were first trapped on a C18 precolumn (300 µm inner diameter x 5 mm; Dionex) at 20 µl/min or 2.5 µL/min with buffer A (2/98 MeCN/H2O in 0.1% formic acid). After 3 min or 4 min of desalting, the precolumn was switched on line with the analytical C18 column (75 µm inner diameter x 50 cm; nanoViper Acclaim PepMapTM RSLC, 2 µm, 100Å, Thermo Scientific) equilibrated in buffer A. Separation was then performed with a linear gradient of 5% to 25% or 30% buffer B (100% MeCN in 0.1% formic acid) at a flow rate of 300 nL/min over 100 min or 91 min. MS full scans were performed in the ultrahigh-field Orbitrap mass analyzer in ranges m/z 400–1500 or m/z 375-1500 with a resolution of 120 000 at m/z 200, ions from each full scan were HCD fragmented and analyzed in the linear ion trap or orbitrap.

For identification, the data were merged and searched against the *S. frugiperda* corn strain proteome [46] using Sequest$^{HF}$ through proteome discoverer (version 2.2) with the *S. frugiperda* CENP-T, CENP-I, CENP-N and Nsl1 candidate added manually to the proteome database. For identification of proteins in the *B. mori* IPs, the data were searched against the *B. mori* proteome. Enzyme specificity was set to trypsin and a maximum of two-missed cleavage sites were allowed. Oxidized methionine, Carbamidomethyl cysteines and N-terminal acetylation were set as variable modifications. Maximum allowed mass deviation was set to 10 ppm for monoisotopic precursor ions and 0.6 Da for MS/MS peaks. The resulting files were further processed using myProMS [66] v3.6. FDR calculation used Percolator and was set to 1% at the peptide level for the whole study.

For the kinetochore IPs on full-length lepidopteran proteins identified proteins that were at least four-fold enriched over the control (the two controls were combined) with a minimum of seven derived peptides were analyzed further (Table S1). For the *S. frugiperda* CENP-T-HFD-FLAG IP (Figure S4) proteins that were at least four-fold enriched over the control with at least 5 derived peptides were selected for further analyses (see above). For Figure 1, known kinetochore homologs or proteins with at least seven derived peptides and that were enriched at least four-fold in three kinetochore immunoprecipitates are listed.

**Immunofluorescence**

Cells were grown on glass coverslips and fixed with ice cold MeOH (anti-CENP-T and anti-Spc24/25), ice cold acetone (anti-Dsn1) and 4% PFA (anti-tubulin), followed by permeabilization using 0.3% Triton X-100 in PBS and blocked in 3% BSA-PBS. The following antibodies were used: rabbit polyclonal anti-CENP-T (rabbit 045), rabbit polyclonal anti-Spc24/25 (rabbit 1621016) and polyclonal rabbit anti-Dsn1 (rabbit 1615031) generated by Covalab (Villeurbanne, FR) at the dilution 1:1000, anti-α-tubulin monoclonal Alexa Fluor 488 (53-4502-80, eBioScience) at 1:1000, anti-FLAG M2 mouse monoclonal (F1804-1MG, Sigma) at 1:1000, anti-phospho Histone H3-Ser10 rat monoclonal (MABE939, Sigma) at 1:1000. For fluorescent conjugated secondary antibodies, we used goat anti-rabbit IgG Alexa Fluor 568 (A-11011, Invitrogen) at 1:1000, goat anti-rat IgG Alexa Fluor 568 (A-11077, Invitrogen) at 1:1000, goat anti-rat IgG Alexa Fluor 488 (A-11006, Invitrogen) at 1:1000, goat anti-mouse IgG Alexa Fluor 488 (A-11029, Invitrogen) at 1:1000, goat anti-mouse IgG Alexa Fluor 568 (A-11004, Invitrogen) at 1:1000 and goat anti-rat IgG Alexa Fluor 633 (A-21094, Invitrogen) at 1:1000. DNA was stained with DAPI (D9542-10MG, Sigma) and samples were mounted in Vectashield Antifade Mounting Medium (H-1000, Vector Labs).

For anti-tubulin staining cells were fixed three and five days after RNAi-mediated depletion using a protocol for the preservation of the whole cytoskelon [68]. Cells were washed with PBS for 5 minutes, then incubated for 10 min at room temperature in 1 mM dithiobis(succinimidyl propionate, DSP) (22585, Thermo Fisher Scientific) in Hank's balanced salt solution (HBSS) (14025050, Gibco), followed by an incubation for 10 min at room temperature in 1 mM DSP in microtubule-stabilizing buffer (MTSB). Cells were next washed for 5 min in 0.5% Triton X-100 in MTSB and then fixed in 4% PFA in MTSB for 15 min at room temperature. After a 5 min wash in PBS, cells were incubated for 5 min in 100 mM glycine in PBS, then washed again in PBS for 5 minutes and finally nuclei were stained with DAPI (D9542-10MG, Sigma) and samples were mounted in Vectashield Antifade Mounting Medium (H-1000, Vector Labs).

**Microscopy**

Images were acquired on Zeiss Axiovert Z1 light microscope. Z-sections were acquired at 0.2μm steps using 100X 1.4 NA oil objective.

Quantification of fluorescence intensity was performed using the Image J software (NIH) on unprocessed TIFF images. Mitotic cells (H3S10ph positive) were quantified. For RNAi depleted cells, we first annotated the cells using an automated system (kind gift from Solène Hervé, Fachinetti lab, UMR144, Institut Curie, Paris, France). H3S10p signals were then used as markers to manually select, using the freehand selections tool, the nuclear area. The mean fluorescence intensity of each nucleus was measured and corrected for background. For background correction, the average the mean intensities of three random circular regions of fixed size (30x30 pixels) placed outside nuclear areas was determined.

To quantify the fluorescent signal intensities for LacO/LacI tethering assays, the mean fluorescence intensities of CENP-T, Dsn1 and Spc24/25 signals were first measured in circular regions of fixed size (10x10 pixels) overlapping LacI foci (visualized by the GFP signals). Then, the mean fluorescence intensity of CENP-T, Dsn1 and Spc24/25 at endogenous loci was determined as the average in three random circular regions of fixed size (10x10 pixels) placed the nuclear area. As before, the background intensity was also measured as the average of mean fluorescence intensities of three random circular regions of fixed size (10x10 pixels). Both the mean fluorescence intensities overlapping the LacI foci or the endogenous loci were corrected with this background value.

For statistical analysis the Mann-Whitney test (unpaired, non-parametric test) was used to compare two ranks, using GraphPad Prism version 8.12 for Mac (GraphPad Software, https://www.graphpad.com). Differences were considered statistically significant at values of P values <0.05.

**RNAi-mediated knock-down**

BmN4-Sid1 cells were grown on coverslips and incubated with 400pg/µl dsRNA for three days. After three days, the medium was change to add another 400pg/µl dsRNA. After three or five days, cells were fixed and processed for IF as described. The following primer fused to T7 promoter sequences were used to generate the DNA templates for dsRNA generation:

GFP_T7_for

TAATACGACTCACTATAGGGAGA GATGCCACCTACGGCAAG

GFP_T7_rev

TAATACGACTCACTATAGGGAGA CGCGGGTCTTGTAGTTGC

BomCenpM_T7_for

TAATACGACTCACTATAGGGAGA TGAATGTTGAAGTAATCGAAAAGG

BomCenpM_T7_rev

TAATACGACTCACTATAGGGAGA TTGCCATAGCATTCACAGGT

BomCenpI_T7_for

TAATACGACTCACTATAGGGAGA CAGTGGATTATGCTATTCAGTGG

BomCenpI_T7_rev

TAATACGACTCACTATAGGGAGA ATTTTCCGGGACACACTCAG

BomCenpN_T7_for

TAATACGACTCACTATAGGGAGA GCCTCTTCAACAATGCTGTG

BomCenpN_T7_rev

TAATACGACTCACTATAGGGAGA AGAGCATCGACACAGGCTTT

BomCenpT_T7_for

TAATACGACTCACTATAGGGAGA GATCCTCCACAAAACCAACC

BomCenpT_T7_rev

TAATACGACTCACTATAGGGAGA TTCTCGCATACCATTTCGTG

BomMis12_T7_for

TAATACGACTCACTATAGGGAGA GGGAACGGATGAGGAATATG

BomMis12_T7_rev

TAATACGACTCACTATAGGGAGA AGCAATGCAACTTCGTCTTTT

BomNsl1_T7_for

TAATACGACTCACTATAGGGAGA GGAGACGAAATGCGAGAATC

BomNsl1_T7_rev

TAATACGACTCACTATAGGGAGA CAGTTTGCCGCCAATTTTAT

BomDsn1_T7_for

TAATACGACTCACTATAGGGAGA CCATCAGTGAAAATGAAATACAACA

BomDsn1_T7_rev

TAATACGACTCACTATAGGGAGA CAAGTGCCATAACTTCTTTGACA

BomSpc24_T7_for

TAATACGACTCACTATAGGGAGA AGATTGGTGTGCCGTGCTAATT

BomSpc24_T7_rev

TAATACGACTCACTATAGGGAGA GTCGGCAGAGTCCACTTCGAAA

BomSpc25_T7_for

TAATACGACTCACTATAGGGAGA CTCATGAAGCCTATTTGCTAACT

BomSpc25_T7_rev

TAATACGACTCACTATAGGGAGA TACTTTATTTTGTTTAATGTTGAGAAA

**RNA blot analyses**

Total RNA was isolated using Trizol following the manufactor's instruction. RNA blots were performed using around 10 µg total RNA (CENP-T, CENP-I, CENP-N, Nsl1, Mis12, Spc25) or polyA-selected mRNAs from 20 µg total RNA (CENP-M, Spc24) per lane. RNA samples from cells incubated for five days with respective dsRNAs were treated with glyoxal using NorthernMax-Gly Sample Loading Dye (Ambion). The samples were loaded on a 1% agarose gel prepared using NorthernMax-Gly Gel Running Buffer (Ambion) according to the manufacturer's instructions. The RNA was blotted onto a Nytran membrane in 20X SSC (175.3 g NaCl, 88.2 g sodium citrate in 1.0 l water adjusted to pH 7) using the TurboBlotter System (Whatman). After UV crosslinking RNA to the membrane, glyoxal treatment was reversed by incubating the membrane in 10 mM Tris-HCl pH 8 for 20 minutes at room temperature. The membrane was incubated in 12 ml QuickHyb solution (Stratagene) with 1.0 mg salmon-sperm ssDNA (Sigma) for 1 hour at 65°C. Body-labeled antisense riboprobes against kinetochore mRNAs and the loading control were prepared by using PCR products as templates for *in vitro* transcription (MaxiScript kit, Ambion). A radiolabeled probe against *B. mori* Rpl32 mRNA served as loading control. After an overnight hybridization at 65°C with radio-labelled probes, the membrane was washed twice in 2X SSC, 0.1% SDS for 5minutes and once in 0.2X SSC, 0.1% SDS for 30 minutes. The membranes were exposed to phosphoimaging plates and analyzed using a Typhoon TRIO Imager.

**CRISPR-mediated genome editing in *B. mori* N4 strain**

The non-diapause strain N4 maintained at the University of Tokyo was used. All larvae were fed with fresh mulberry leaves or artificial diet SilkMate (NOSAN) under a continuous cycle of 12-h light and 12-h darkness at 25°C. Unique single-guide RNA (sgRNA) target sequences in the silkworm genome were selected using CRISPRdirect (https://crispr.dbcls.jp/) [69]. The sequence specificity in N4 strain was also checked by SilkBase (http://silkbase.ab.a.u-tokyo.ac.jp). Primers used for sgRNA transcription *in vitro* are as follows: forward primer sequence: GAAATTAATACGACTCACTATA gatgaaccagaaaacagtgcGTTTTAGAGCTAGAAATAGC; reverse primer sequence: AAAAGCACCGACTCGGTGCCACTTTTTCAAGTTGATAACGGACTAGCCTTATTTTAACTTGCTATTTCTAGCTCT AAAAC. The sgRNA was transcribed *in vitro* according to a method reported previously [70]. A mixture of sgRNA (400 ng/µL) and Cas9 Nuclease protein NLS (120 ng/µL; NIPPON GENE) in injection buffer (100 mM KOAc, 2 mM $Mg(OAc)_2$, 30 mM HEPES-KOH; pH 7.4) was injected into each egg within 3 h after oviposition [71]. The injected embryos were incubated at 25°C in a humidified Petri dish until hatching. Injected individuals were crossed with non-injected individuals to obtain $G_1$ broods. To identify $G_1$ moths in which mutant alleles were transmitted from $G_0$, genomic DNA was extracted from a $G_1$ adult leg using the hot sodium hydroxide and Tris (HotSHOT) method [72]. Genomic PCR was performed using KOD One™ (TOYOBO) under the following conditions: 35 cycles of denaturation at 98°C for 10 s, annealing at 60°C for 5 s, extension at 68°C for 5 s. Primers used for mutation screening are as follows: forward primer sequence: ATCCAACGCAAGTAATGACGAT; reverse primer sequence: ATCAGTGTTCGGGCTATTATCA. The PCR products were denatured and reannealed at 95°C for 10 min, followed by gradual cooling to 25°C. Mutations at the target site were detected by heteroduplex mobility assay using the MultiNA microchip electrophoresis system (SHIMADZU) with the DNA-500 reagent kit [73,74]. A mutation at the target site was sequenced using BigDye® Terminator v3.1 Cycle Sequencing Kit (Applied Biosystems) and ABI PRISM® 3130xl Genetic Analyzer (Applied Biosystems). We maintained the mutant line and obtained eggs carrying the homozygous mutation by crossing between two heterozygous mutants.

Figure S1

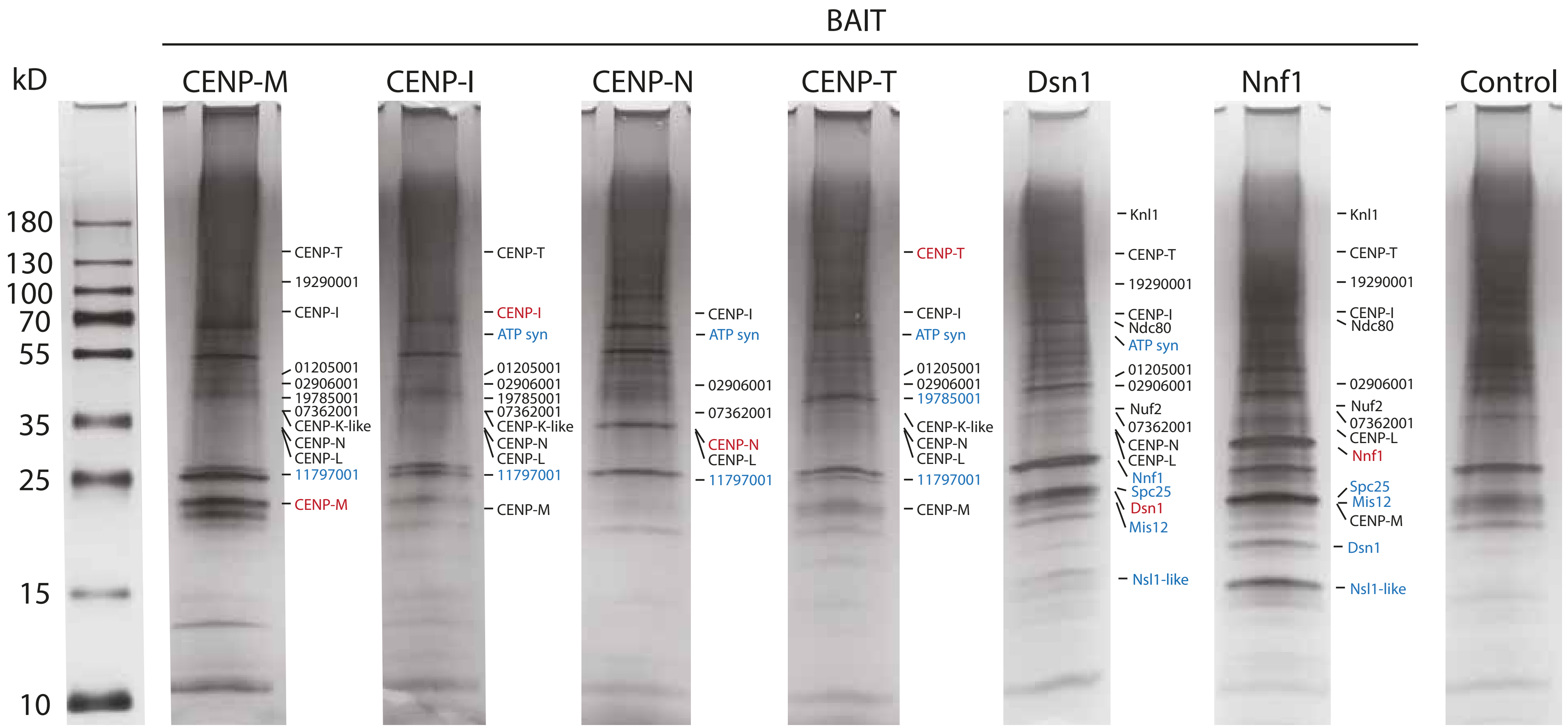
BAIT
kD
CENP-M
CENP-I
CENP-N
CENP-T
Dsn1
Nnf1
Control
180
130
100
70
55
35
25
15
10
CENP-T
19290001
CENP-I
01205001
02906001
19785001
07362001
CENP-K-like
CENP-N
CENP-L
11797001
CENP-M
CENP-T
CENP-I
ATP syn
01205001
02906001
19785001
07362001
CENP-K-like
CENP-N
CENP-L
11797001
CENP-M
CENP-I
ATP syn
02906001
07362001
CENP-N
CENP-L
11797001
CENP-T
CENP-I
ATP syn
01205001
02906001
19785001
CENP-K-like
CENP-N
CENP-L
11797001
CENP-M
Knl1
CENP-T
19290001
CENP-I
Ndc80
ATP syn
01205001
02906001
Nuf2
07362001
CENP-N
CENP-L
Nnf1
Spc25
Dsn1
Mis12
Nsl1-like
Knl1
CENP-T
19290001
CENP-I
Ndc80
02906001
Nuf2
07362001
CENP-L
Nnf1
Spc25
Mis12
CENP-M
Dsn1
Nsl1-like

Figure S2

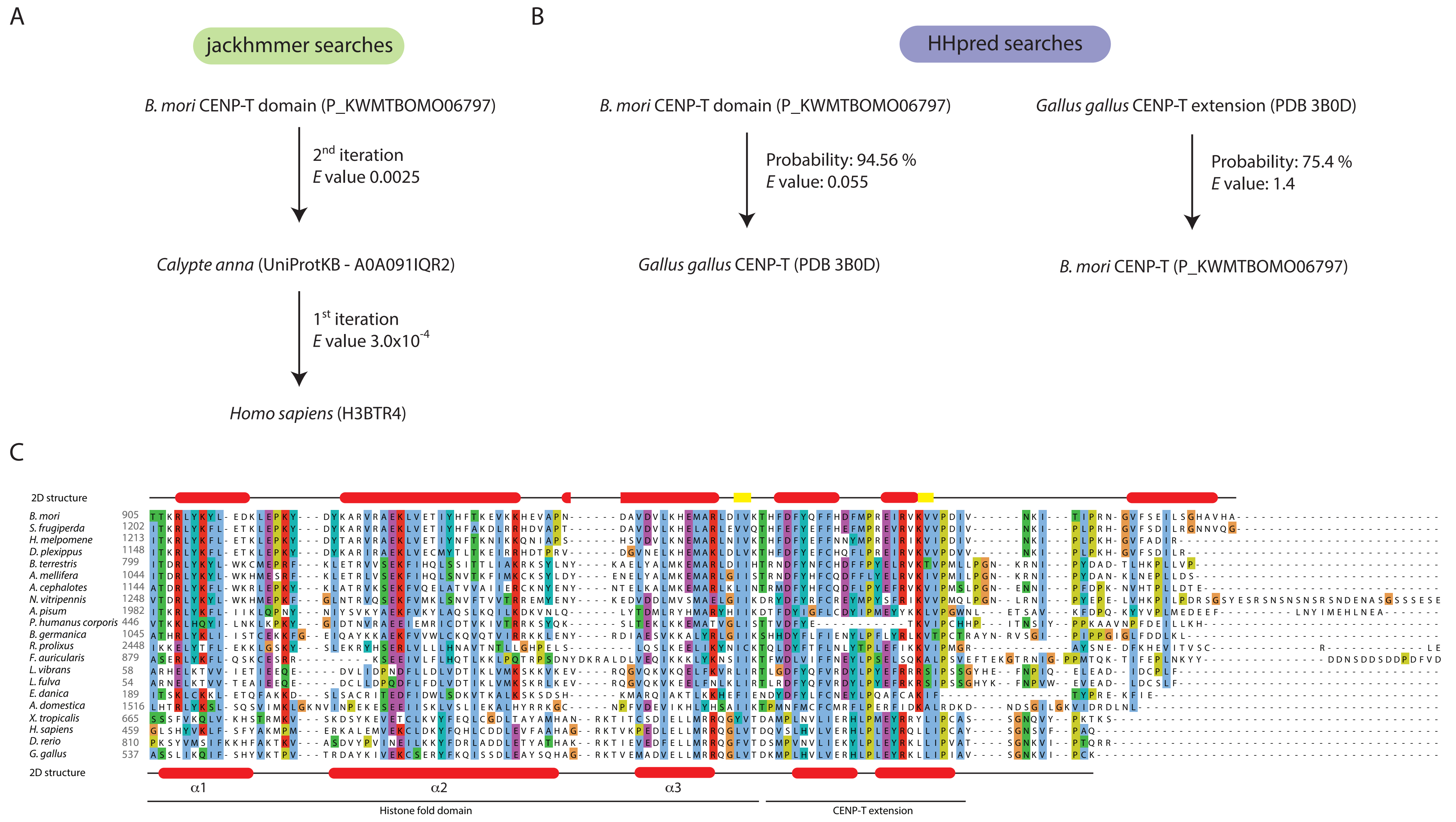

A
jackhmmer searches
B. mori CENP-T domain (P_KWMTBOMO06797)
2nd iteration
E value 0.0025
Calypte anna (UniProtKB - A0A091IQR2)
1st iteration
E value 3.0x10-4
Homo sapiens (H3BTR4)
B
HHpred searches
B. mori CENP-T domain (P_KWMTBOMO06797)
Probability: 94.56 %
E value: 0.055
Gallus gallus CENP-T (PDB 3B0D)
Gallus gallus CENP-T extension (PDB 3B0D)
Probability: 75.4 %
E value: 1.4
B. mori CENP-T (P_KWMTBOMO06797)
C
2D structure
B. mori 905
S. frugiperda 1202
H. melpomene 1213
D. plexippus 1148
B. terrestris 799
A. mellifera 1044
A. cephalotes 1144
N. vitripennis 1248
A. pisum 1982
P. humanus corporis 446
B. germanica 1045
R. prolixus 2448
F. auricularis 879
L. vibrans 58
L. fulva 54
E. danica 189
A. domestica 1516
X. tropicalis 665
H. sapiens 459
D. rerio 810
G. gallus 537
2D structure
α1
α2
α3
Histone fold domain
CENP-T extension

Figure S3

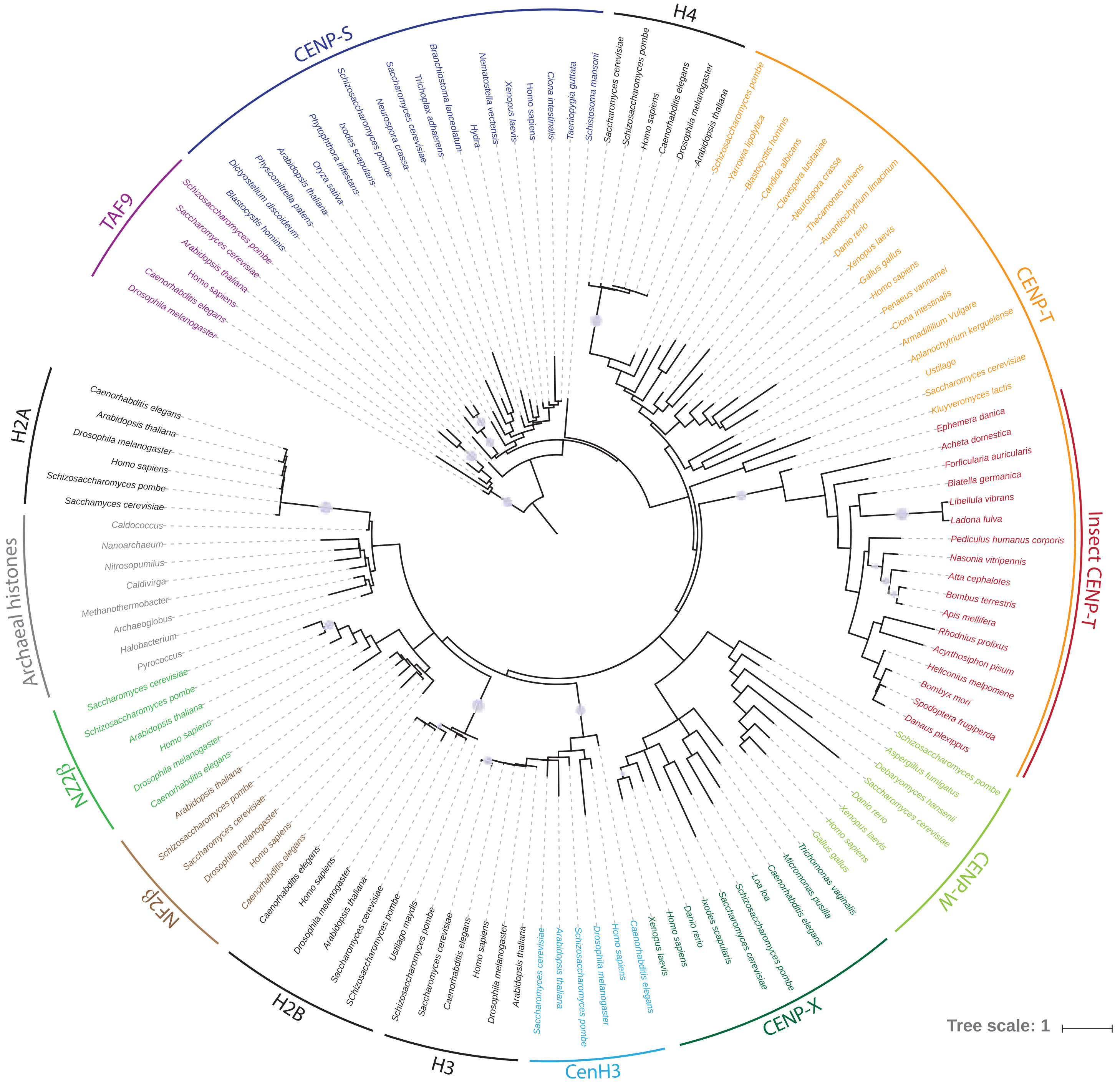

H4
Saccharomyces cerevisiae
Schizosaccharomyces pombe
Homo sapiens
Caenorhabditis elegans
Drosophila melanogaster
Arabidopsis thaliana
CENP-T
Schizosaccharomyces pombe
Yarrowia lipolytica
Blastocystis hominis
Candida albicans
Clavispora lusitaniae
Neurospora crassa
Thecamonas trahens
Aurantiochytrium limacinum
Danio rerio
Xenopus laevis
Gallus gallus
Homo sapiens
Penaeus vannamei
Ciona intestinalis
Armadillilium Vulgare
Aplanochytrium kerguelense
Ustilago
Saccharomyces cerevisiae
Kluyveromyces lactis
Insect CENP-T
Ephemera danica
Acheta domestica
Forficularia auricularis
Blatella germanica
Libellula vibrans
Ladona fulva
Pediculus humanus corporis
Nasonia vitripennis
Atta cephalotes
Bombus terrestris
Apis mellifera
Rhodnius prolixus
Acyrthosiphon pisum
Heliconius melpomene
Bombyx mori
Spodoptera frugiperda
Danaus plexippus
CENP-W
Schizosaccharomyces pombe
Aspergillus fumigatus
Debaryomyces hansenii
Saccharomyces cerevisiae
Danio rerio
Xenopus laevis
Homo sapiens
Gallus gallus
Trichomonas vaginalis
Micromonas pusilla
Caenorhabditis elegans
CENP-X
Loa loa
Schizosaccharomyces pombe
Saccharomyces cerevisiae
Ixodes scapularis
Danio rerio
Homo sapiens
Xenopus laevis
CenH3
Caenorhabditis elegans
Homo sapiens
Drosophila melanogaster
Schizosaccharomyces pombe
Arabidopsis thaliana
Saccharomyces cerevisiae
H3
Arabidopsis thaliana
Drosophila melanogaster
Homo sapiens
Caenorhabditis elegans
Saccharomyces cerevisiae
Schizosaccharomyces pombe
Ustilago maydis
H2B
SChizosaccharomyces pombe
Saccharomyces cerevisiae
Arabidopsis thaliana
Drosophila melanogaster
Homo sapiens
Caenorhabditis elegans
NF2B
Caenorhabditis elegans
Homo sapiens
Drosophila melanogaster
Saccharomyces cerevisiae
Schizosaccharomyces pombe
Arabidopsis thaliana
NZ2B
Caenorhabditis elegans
Drosophila melanogaster
Homo sapiens
Arabidopsis thaliana
Schizosaccharomyces pombe
Saccharomyces cerevisiae
Archaeal histones
Pyrococcus
Halobacterium
Archaeoglobus
Methanothermobacter
Caldivirga
Nitrosopumilus
Nanoarchaeum
Caldococcus
H2A
Sacchamyces cerevisiae
Schizosaccharomyces pombe
Homo sapiens
Drosophila melanogaster
Arabidopsis thaliana
Caenorhabditis elegans
TAF9
Drosophila melanogaster
Caenorhabditis elegans
Homo sapiens
Arabidopsis thaliana
Saccharomyces cerevisiae
Schizosaccharomyces pombe
CENP-S
Blastocystis hominis
Dictyostelium discoideum
Physcomitrella patens
Arabidopsis thaliana
Oryza sativa
Phytophthora infestans
Ixodes scapularis
Schizosaccharomyces pombe
Neurospora crassa
Saccharomyces cerevisiae
Trichoplax adhaerens
Branchiostoma lanceolatum
Hydra
Nematostella vectensis
Xenopus laevis
Homo sapiens
Ciona intestinalis
Taeniopygia guttata
Schistosoma mansoni
Tree scale: 1

### Figure S4

| *S. frugiperda* ID | *B. mori* ID | Number of peptides in IP | Coverage (%) | Number of peptides in control | MWG (kDa) | Description |
|---|---|---|---|---|---|---|
| GSSPFG00032913001 | KWMTBOMO14835 | 51 | 30.4 | 6 | 105.2 | Phosphatidylinositol 3-kinase 3 isoform X1 |
| This study | KWMTBOMO06797 | 48 | 12.1 | 1 | 150.0 | CENP-T |
| GSSPFG00001205001 | KWMTBOMO14835 | 32 | 44.7 | 3 | 51.3 | Coil-coiled RWD like protein |
| This study | KWMTBOMO02221 | 28 | 36.4 | 0 | 76.0 | CENP-I |
| GSSPFG00011797001 | KWMTBOMO06154 | 20 | 66.0 | 0 | 28.0 | unknown |
| GSSPFG00019785001 | KWMTBOMO09290 | 18 | 34.2 | 0 | 43.9 | Coil-coiled RWD like protein |
| GSSPFG00011349001 | LOC105842400 | 13 | 74.1 | 0 | 12.2 | Annotated as non-coding RNA |
| GSSPFG00009519001 | KWMTBOMO00944 | 12 | 47.2 | 0 | 8.1 | Uncharacterized protein |
| GSSPFG00023246001 | KWMTBOMO11447 | 10 | 24.5 | 0 | 34.7 | CENP-L |
| GSSPFG00020888001 | KWMTBOMO11666 | 7 | 24.5 | 1 | 55.1 | Transitional endoplasmic reticulum ATPase TER94 |
| GSSPFG00002981001 | KWMTBOMO06965 | 7 | 27.9 | 1 | 59.8 | Uncharacterized protein |
| GSSPFG00005844001 | KWMTBOMO03030 | 6 | 8.6 | 0 | 85.9 | Atlastin |
| GSSPFG00013028001 | KWMTBOMO07723 | 5 | 7.4 | 0 | 129.9 | Eukaryotic translation initiation factor 5B |

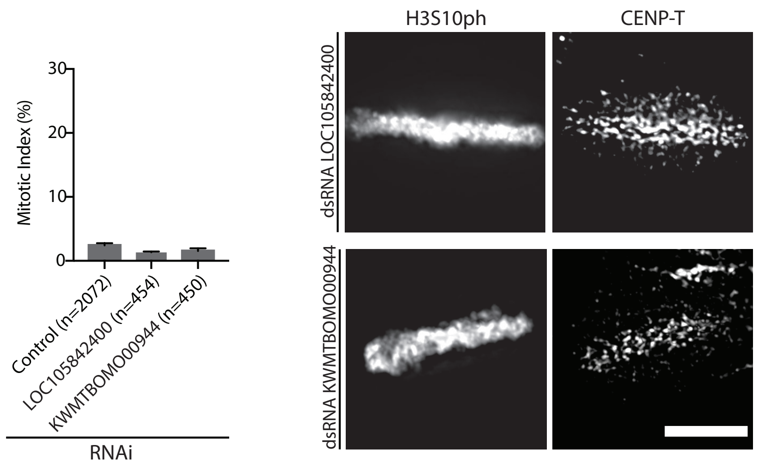

Figure S5

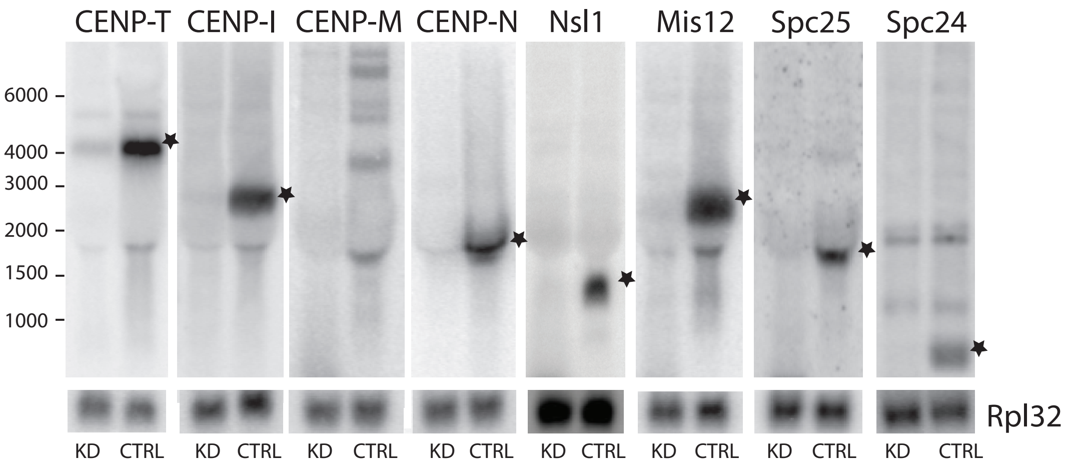

CENP-T
CENP-I
CENP-M
CENP-N
Nsl1
Mis12
Spc25
Spc24
6000
4000
3000
2000
1500
1000
Rpl32
KD
CTRL

Figure S6

A

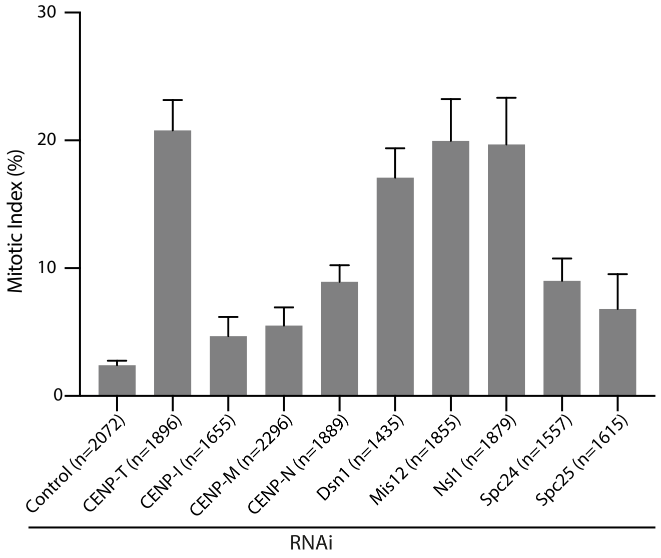
30
20
10
0
Mitotic Index (%)
Control (n=2072)
CENP-T (n=1896)
CENP-I (n=1655)
CENP-M (n=2296)
CENP-N (n=1889)
Dsn1 (n=1435)
Mis12 (n=1855)
Nsl1 (n=1879)
Spc24 (n=1557)
Spc25 (n=1615)
RNAi

B

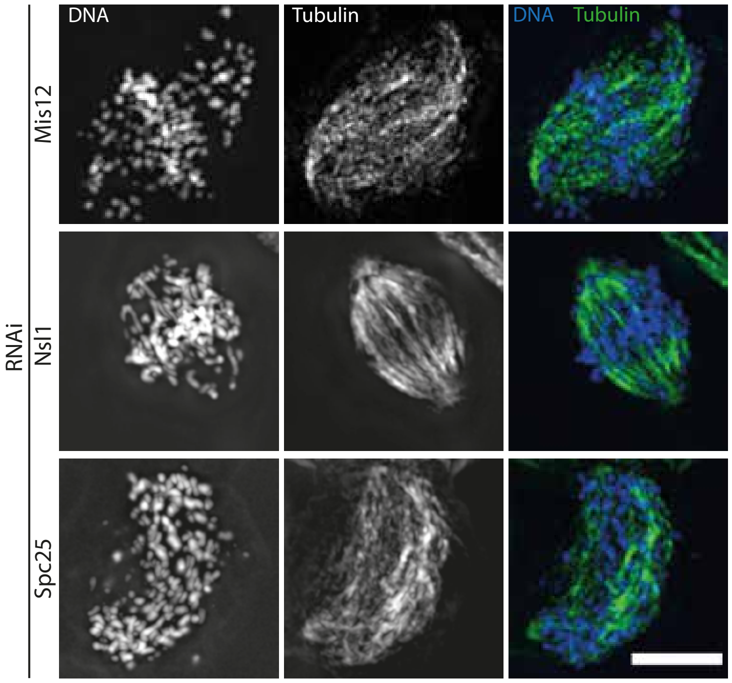
DNA
Tubulin
DNA Tubulin
Mis12
Nsl1
Spc25
RNAi

Figure S7

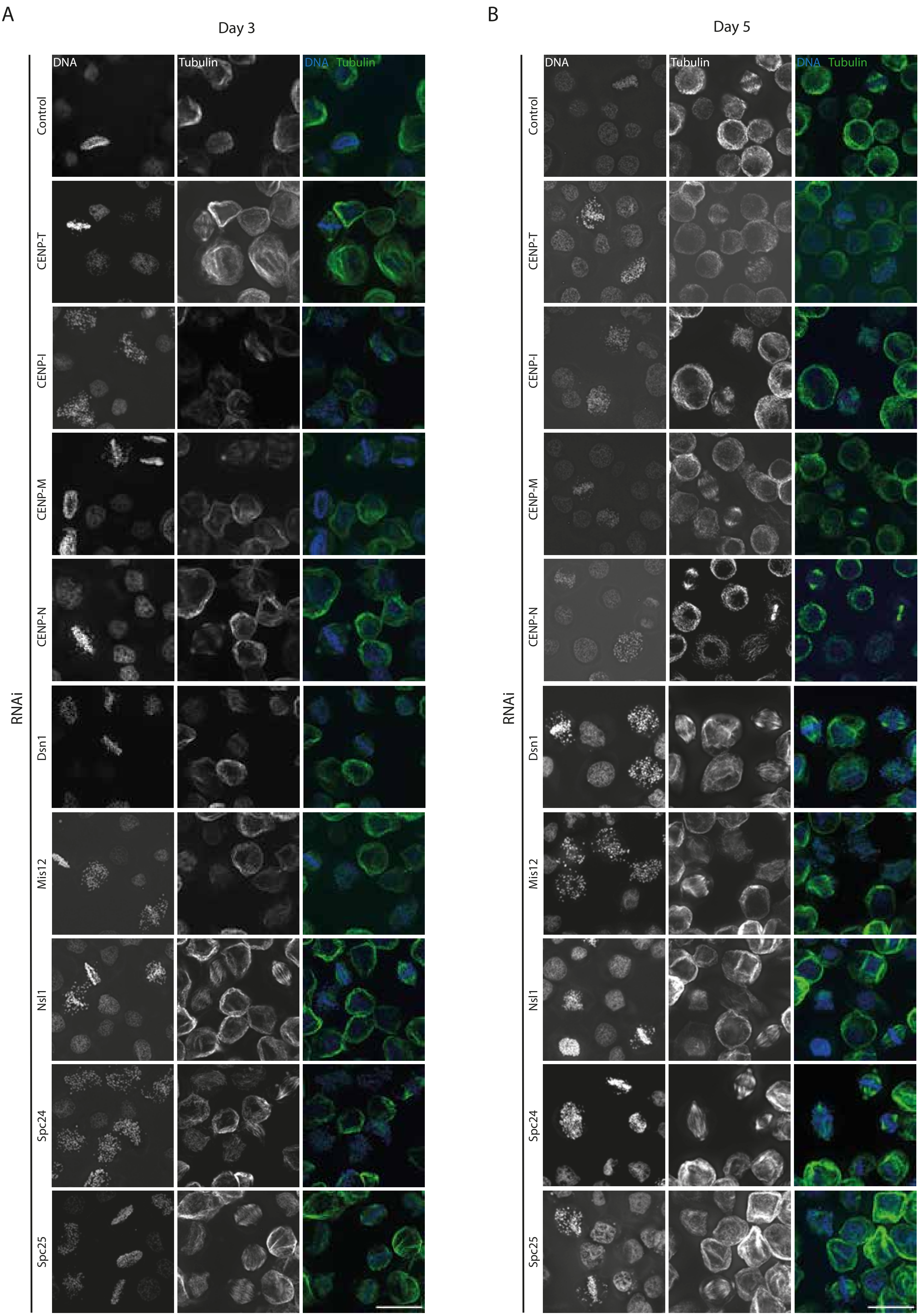
A
Day 3
DNA
Tubulin
DNA Tubulin
Control
CENP-T
CENP-I
CENP-M
CENP-N
Dsn1
Mis12
Nsl1
Spc24
Spc25
RNAi
B
Day 5
DNA
Tubulin
DNA Tubulin
Control
CENP-T
CENP-I
CENP-M
CENP-N
Dsn1
Mis12
Nsl1
Spc24
Spc25
RNAi

Figure S8

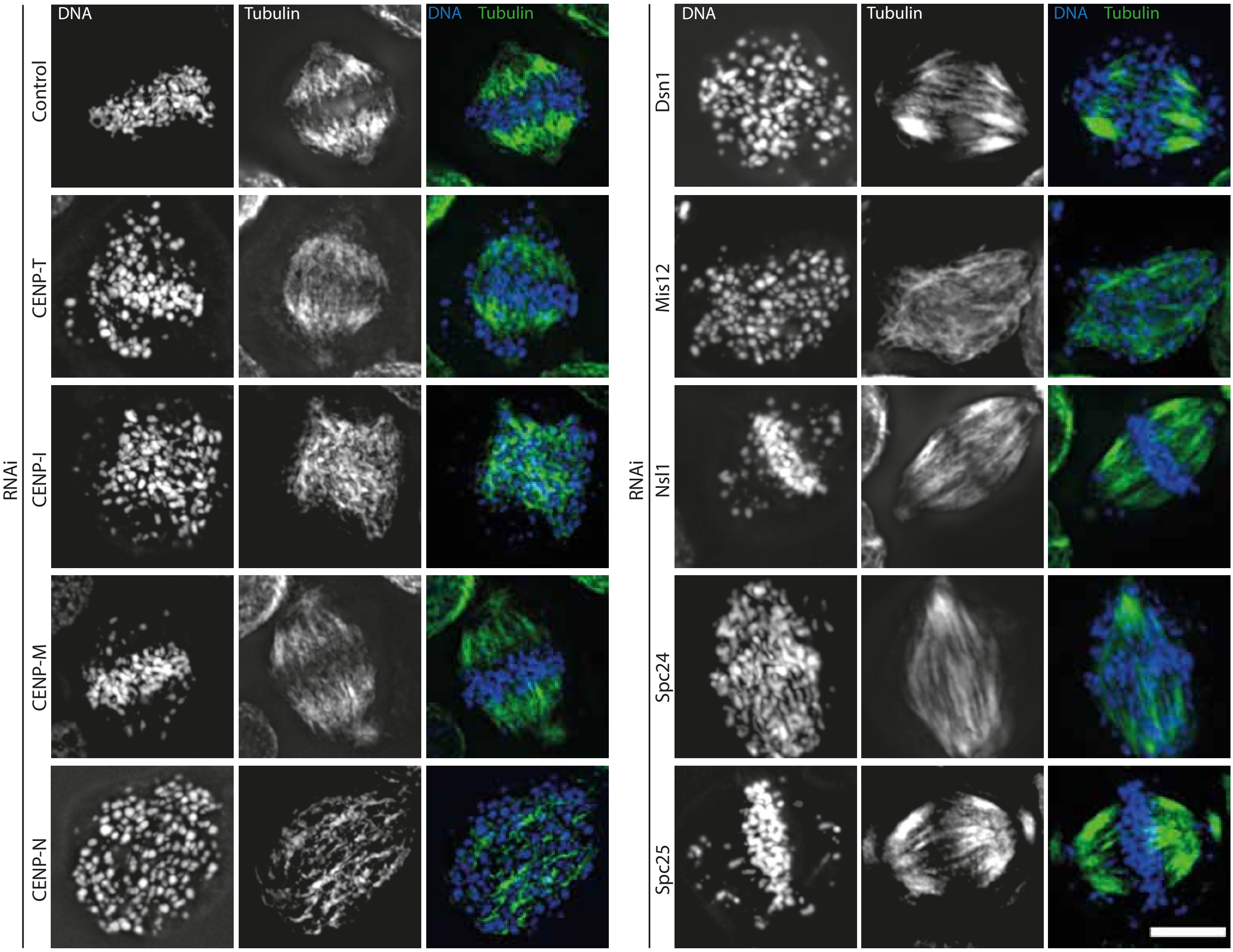

DNA
Tubulin
DNA Tubulin
RNAi
Control
CENP-T
CENP-I
CENP-M
CENP-N
Dsn1
Mis12
Nsl1
Spc24
Spc25

Figure S9

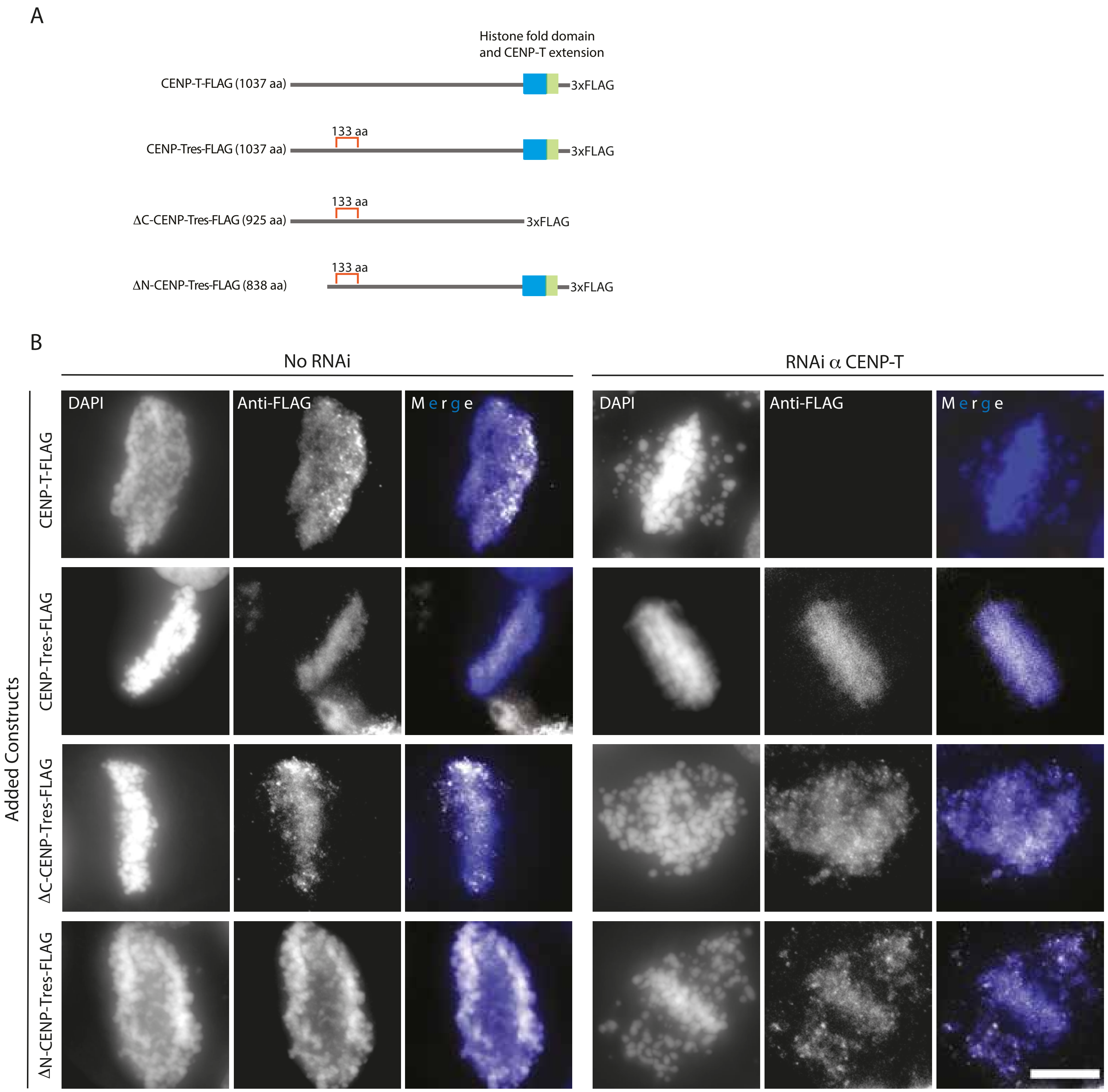
A
Histone fold domain
and CENP-T extension
CENP-T-FLAG (1037 aa)
3xFLAG
133 aa
CENP-Tres-FLAG (1037 aa)
3xFLAG
133 aa
ΔC-CENP-Tres-FLAG (925 aa)
3xFLAG
133 aa
ΔN-CENP-Tres-FLAG (838 aa)
3xFLAG
B
No RNAi
RNAi α CENP-T
DAPI
Anti-FLAG
Merge
DAPI
Anti-FLAG
Merge
Added Constructs
CENP-T-FLAG
CENP-Tres-FLAG
ΔC-CENP-Tres-FLAG
ΔN-CENP-Tres-FLAG

Figure S10

A

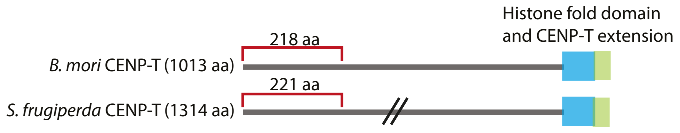


| *B. mori* ID | Number of peptides | Coverage (%) | MWG (kDa) | Description |
|---|---|---|---|---|
| P_KWMTBOMO06797 | 57 | 51.0 | 116.6 | CENP-T |
| P_KWMTBOMO05194 | 10 | 16.5 | 81 | CD2-associated protein |
| P_KWMTBOMO14073 | 10 | 9.1 | 171.2 | Acetyl-CoA carboxylase isoform X2 |
| P_KWMTBOMO06603 | 6 | 10.0 | 104.4 | Uncharacterized protein LOC101742952 |
| P_KWMTBOMO14420 | 6 | 20.1 | 106.6 | Heterogeneous nuclear ribonucleoprotein U-like protein 1 |

B

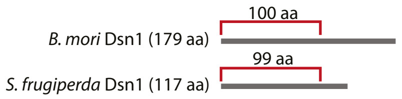


| *B. mori* ID | Number of peptides | Coverage (%) | MWG (kDa) | Description |
|---|---|---|---|---|
| P_KWMTBOMO11045 | 32 | 18.5 | 268.5 | Spectrin beta chain |
| P_KWMTBOMO08735 | 17 | 69.8 | 21.1 | Dsn1 |
| P_KWMTBOMO11745 | 13 | 41.6 | 37.7 | DNA repair protein RAD52 homolog |
| P_KWMTBOMO08648 | 11 | 6.6 | 273.4 | Spectrin alpha chain |
| P_KWMTBOMO06463 | 7 | 6.1 | 175.3 | DNA topoisomerase 2 |
| P_KWMTBOMO14420 | 7 | 9.9 | 106.6 | Heterogeneous nuclear ribonucleoprotein U-like protein 1 |

C

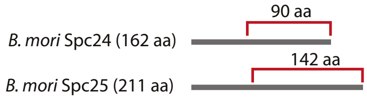


IP of His-Spc24 (72-162aa) - Spc25(69-142aa) from baculoviral-infected SF9 cells

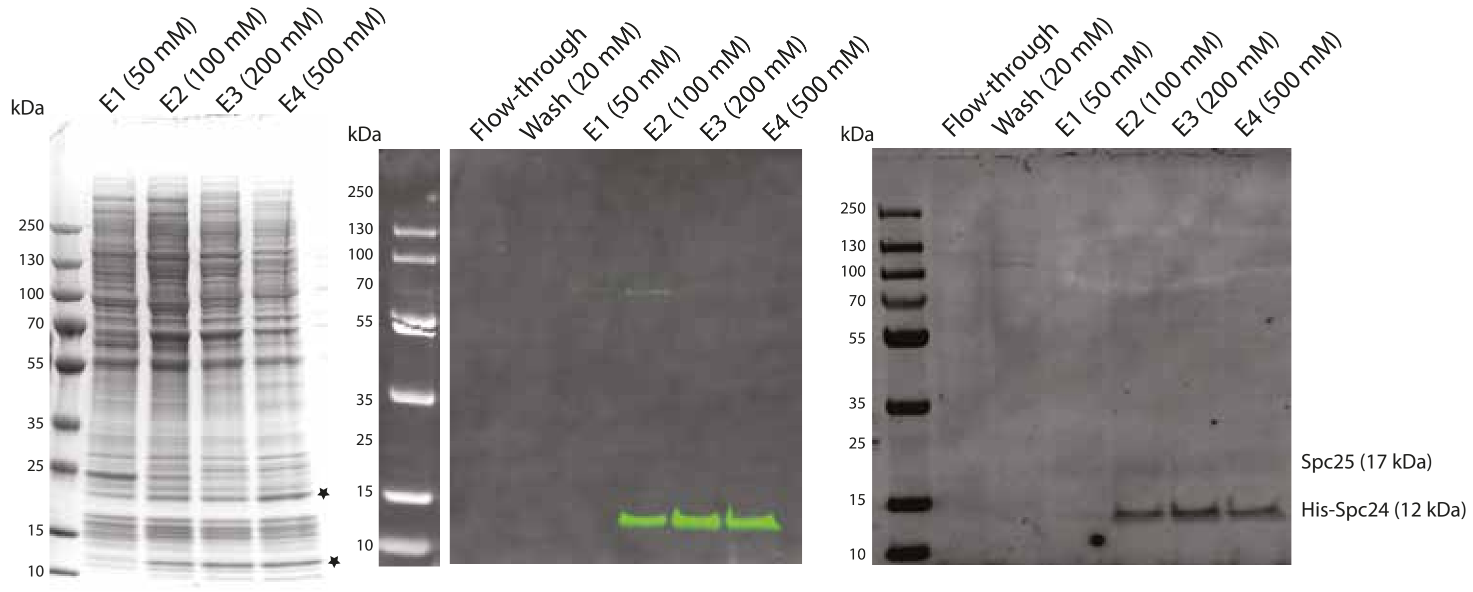


Commassi

Western Blot α-His

Western Blot α-Spc24/Spc25

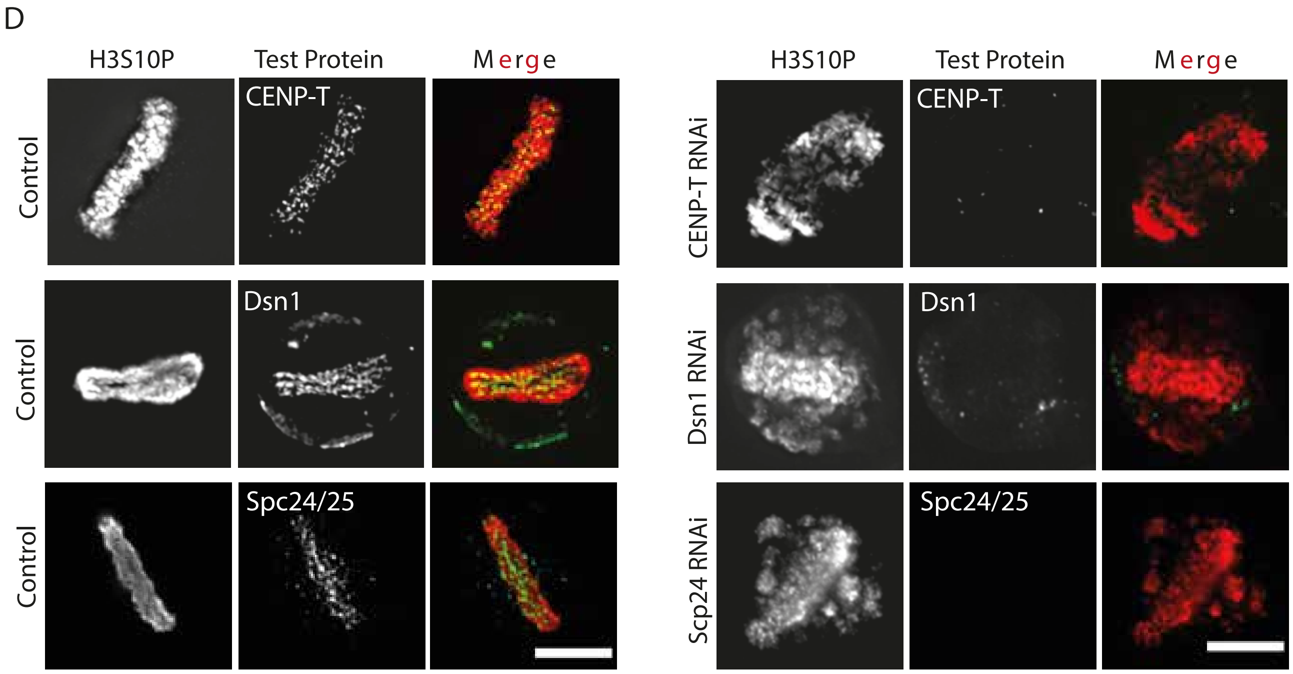
D
H3S10P
Test Protein
Merge
H3S10P
Test Protein
Merge
Control
CENP-T
CENP-T RNAi
CENP-T
Control
Dsn1
Dsn1 RNAi
Dsn1
Control
Spc24/25
Scp24 RNAi
Spc24/25

Figure S11

A

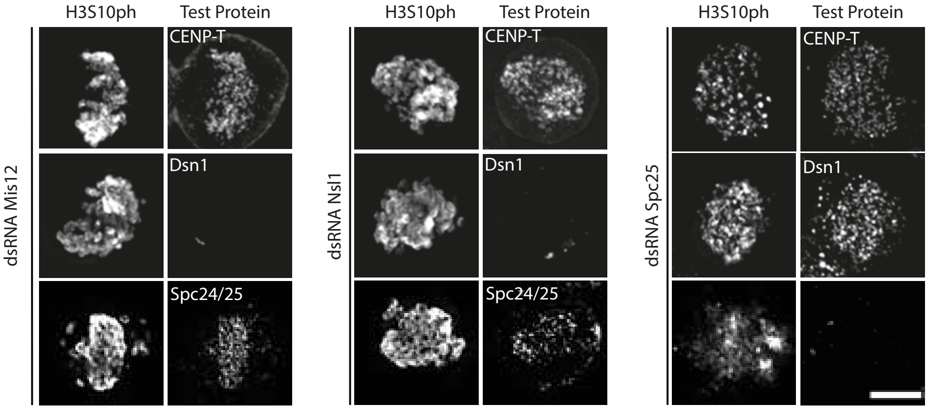


B

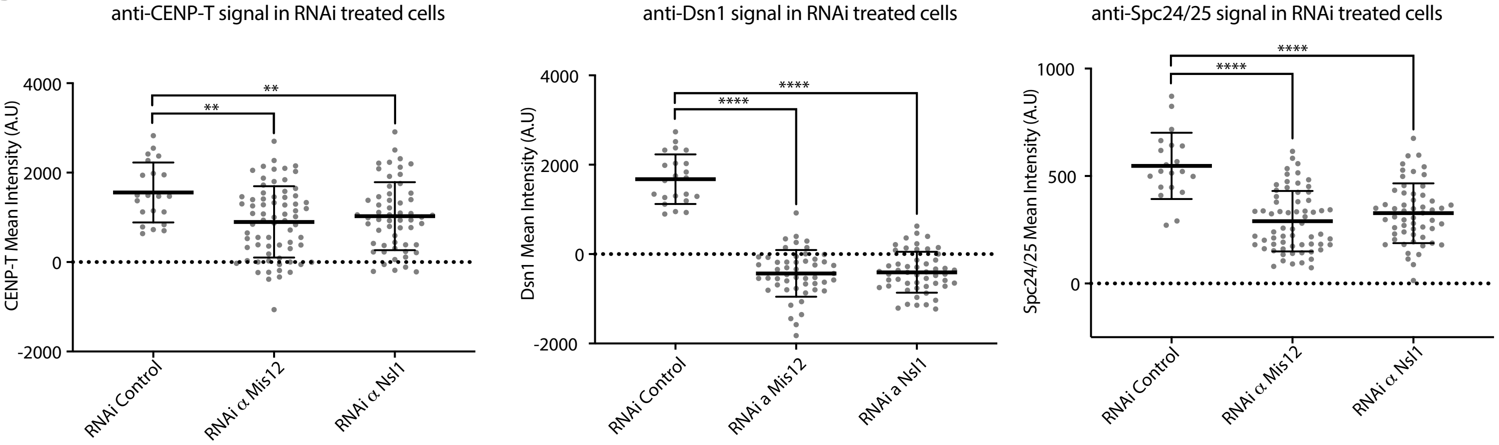

**Table S3: Hatchability of CENP-T mutant strain**

| Crossing pattern (♀ × ♂) | No. of hatched eggs (%) | No. of unhatched eggs (%) | No. of eggs died just before hatching (%) | Total |
|---|---|---|---|---|
| | 174 (88.3) | 7 (3.6) | 16 (8.1) | 197 |
| +/+ × +/+ | 203 (90.2) | 8 (3.6) | 14 (6.2) | 225 |
| | 153 (78.5) | 21 (10.8) | 21 (10.8) | 195 |
| | 227 (87.3) | 15 (5.8) | 18 (6.9) | 260 |
| +/+ × KO/+ | 218 (89.7) | 2 (0.8) | 23 (9.5) | 243 |
| | 178 (83.2) | 14 (6.5) | 22 (10.3) | 214 |
| KO/+ × +/+ | 216 (87.8) | 14 (5.7) | 16 (6.5) | 246 |
| | 227 (90.8) | 10 (4.0) | 13 (5.2) | 250 |
| | 171 (69.5) | 70 (28.5) | 5 (2.0) | 246 |
| KO/+ × KO/+ | 125 (63.5) | 68 (34.5) | 4 (2.0) | 197 |
| | 144 (67.3) | 64 (29.9) | 6 (2.8) | 214 |
| | 184 (69.7) | 74 (28.0) | 6 (2.3) | 264 |

**Table S4 Genotypes of hatched larvae from the cross between two heterozygous CENP-T mutants**

| Crossing pattern (♀ × ♂) | Number of larvae | | | |
|---|---|---|---|---|
| KO/+ × KO/+ | +/+ (%) | KO/+ (%) | KO/KO (%) | Total |
| | 11 (34.4) | 21 (65.6) | 0 (0) | 32 |

**Table S5**

Plasmids generated in this study

| | |
|---|---|
| 143 | *S. frugiperda* CENP-M-FLAG pIBV5 |
| 144 | *S. frugiperda* FLAG-CENP-I pIBV5 |
| 145 | *S. frugiperda* CENP-N-FLAG pIBV5 |
| 142 | *S. frugiperda* CENP-T-FLAG pIBV5 |
| 146 | *S. frugiperda* Dsn1-FLAG pIBV5 |
| 147 | *S. frugiperda* Nnf1-FLAG pIBV5 |
| 150 | *S. frugiperda* CENP-T-HFDextension-FLAG pIBV5 |
| 96 | *B. mori* CENP-T-FLAG pIZV5 |
| 44 | *B. mori* CENP-Tres-FLAG pIZV5 |
| 128 | *B. mori* ΔC-CENP-Tres-FLAG pIZV5 |
| 81 | *B. mori* ΔN-CENP-Tres-FLAG pIZV5 |
| 119 | *S. frugiperda* CENP-T-GFP-LacI pIZV5 |
| 136 | *S. frugiperda* ΔN-CENP-T-GFP-LacI pIZV5 |
| 149 | pVS1-LacO-Blasticidin |
| 99 | LacI-GFP pIBV5 |
| b027 | *S. frugiperda* SUMO-6xHis-CENP-T (1-122) pT7 |
| b030 | *B. mori* SUMO-6xHis-CENP-T (1-218) pT7 |
| b043 | *B. mori* 6xHis-Spc24(73-162)-Spc25(70-211) pRSF-DUET1 |
| b036 | *S. frugiperda* 6xHis Dsn1(1-99) pRSF-DUET1 |
| b035 | *B. mori* 6xHis Dsn1(1-100) pRSF-DUET1 |
| b109 | *B. mori* 6xHis-Spc24(73-162)-Spc25(70-211) pFASTBac-Dual |

**Table S6**
Antibodies generated in this study

| Antibody | Source | Identifier |
|---|---|---|
| Rabbit polyclonal anti-CENP-T | This study | |
| Rabbit polyclonal anti-Spc24/25 | This study | |
| Rabbit polyclonal anti-Dsn1 | This study | |
| Mouse monoclonal anti-6x His tag | Abcam | Ab18184 |
| Goat polyclonal IRDye 680RD anti-Rabbit IgG | LI-COR | 926-68071 |
| Donkey polyclonal IRDye 800CW anti-mouse IgG | LI-COR | 926-32212 |
| Mouse monoclonal anti-3XFLAG M2 beads | Sigma | M8823 |
| Mouse monoclonal anti-FLAG M2 | Sigma | F1804-1MG |
| Rat monoclonal anti-phospho Histone H3 (Ser10), clone 6G8B7 | Sigma | MABE939 |
| Monoclonal anti-α-tubulin Alexa Fluor 488 | eBioScience | 53-4502-80 |
| Polyclonal Goat anti-Rabbit IgG Alexa Fluor 568 | Invitrogen | A-11011 |
| Polyclonal Goat anti-Rat IgG Alexa Fluor 568 | Invitrogen | A-11077 |
| Polyclonal Goat anti-Rat IgG Alexa Fluor 488 | Invitrogen | A-11006 |
| Polyclonal Goat anti-Mouse IgG Alexa Fluor 488 | Invitrogen | A-11029 |
| Polyclonal Goat anti-Mouse IgG Alexa Fluor 568 | Invitrogen | A-11004 |
| Polyclonal Goat anti-Rat IgG Alexa Fluor 633 | Invitrogen | A-21094 |
